# Behavioural stress feedback loops in benthic invertebrates caused by pH drop-induced metabolites

**DOI:** 10.1101/2021.04.16.440165

**Authors:** Lauric Feugere, Lauren Angell, James Fagents, Rebecca Nightingale, Kirsty Rowland, Saffiyah Skinner, Jorg Hardege, Helga Bartels-Hardege, Katharina C. Wollenberg Valero

## Abstract

Studies on pH stress in marine animals typically focus on direct or species-specific aspects. We here test the hypothesis that a drop to pH = 7.6 indirectly affects the intra- and interspecific interactions of benthic invertebrates by means of chemical communication. We recorded fitness-relevant behaviours of small hermit crabs *Diogenes pugilator*, green shore crabs *Carcinus maenas*, and harbour ragworms *Hediste diversicolor* in response to short-term pH drop, and to putative stress metabolites released by conspecifics or gilt-head sea bream *Sparus aurata* during 30 minutes of acute pH drop. Not only did acute pH drop itself impair time to find a food cue in small hermit crabs and burrowing in harbour ragworms, but similar effects were observed under exposure to pH drop-induced stress metabolites. Stress metabolites from *S. aurata,* but not its regular control metabolites, also induced avoidance responses in all recipient species. Here, we confirm that a short-term abrupt pH drop, an abiotic stressor, has the capacity to trigger the release of metabolites which induce behavioural responses in conspecific and heterospecific individuals, which can be interpreted as a behavioural cost. Our findings that stress responses can be indirectly propagated through means of chemical communication warrant further research to confirm the effect size of the behavioural impairments caused by stress metabolites and to characterise their chemical nature.

## Introduction

Compared to the open ocean, coastal areas and particularly intertidal zones are highly variable environments characterised by abrupt changes in water parameters. This includes fluctuations in pH beyond 0.3 units, the levels predicted for average change related to the process of ocean acidification towards the end of the century (Caldeira and Wickett, 2003; Sabine et al., 2004; IPCC, 2019). While this may mean that intertidal species are more resilient to climate change due to their acquired tolerances of pH fluctuations, it also forces them to live more frequently near their physiological tolerance limits (Truchot and Duhamel-Jouve, 1980; Hofmann et al., 2011; Sokolova, 2018; Wolfe et al., 2020). Superimposing ocean acidification on natural pH fluctuations may further increase the variability of organisms’ responses (Eriander et al., 2015). Therefore, organisms inhabiting intertidal areas are interesting models to study the effects of short-term pH fluctuations within the IPCC predicted range. A low environmental pH can alter animal behaviour through several direct or indirect pathways, which include (i) deviation of energy budgets towards the stress response (Pörtner, 2008), (ii) fleeing to avoid the sources of stress (Pörtner and Peck, 2010; Abreu et al., 2016), (iii) disrupted information detection and processing leading to impaired decision-making (Briffa et al., 2012), and (iv) alteration of the chemical signals themselves impacting the sensory environment and the transfer of information (Wyatt et al., 2014; Roggatz et al., 2016, 2019). Behavioural effects triggered by lowered pH are known to occur in different taxonomic groups such as crustaceans (de la Haye et al., 2011), marine ragworms (Bond, 2018), and fish (Munday et al., 2009), although recent research debate on both their ubiquitousness and their effect size (Clark et al., 2020a; Clements et al., 2020b, but see Clark et al., 2020b; Munday et al., 2020).

In aquatic environments, where visibility can be limited, infochemicals and chemosensory functions, such as pheromones used for mating (Karlson and Lüscher, 1959) are crucial for communication (Hardege, 1999; Jordão and Volpato, 2000; Hay, 2009; Ashur et al., 2017). However, they are also a mechanism to propagate stress – but to date mostly researched within the context of biotic stressors. For example, alarm cues released following mechanical damage from fish epidermal club cells can trigger panic reactions in conspecifics and heterospecifics (Toa et al., 2004; Júnior et al., 2010). Such mechanisms of chemical communication also occur in invertebrates, as evidenced by the reduced out-of-burrow activity of the marine polychaete *Alitta virens* exposed to damaged conspecifics (Watson et al., 2005; Ende et al., 2017). Disturbance cues refer to chemicals that may be stored in gill epithelium or urine, and are released voluntarily or involuntarily by disturbed or stressed prey, to induce early antipredator responses in recipients to anticipate potential threats (Bairos-Novak et al., 2017; Goldman et al., 2020a). The central role of chemical communication in the aquatic environment and the recent evidence of its impairment by ocean acidification pinpoints the need for a deeper understanding of such potential environmental modulation of chemical signalling (Chivers et al., 2013).

Overall, though, it is not well understood yet whether abiotic stressors such as pH drop can also induce the release of chemical cues and whether these can propagate the stress response to other individuals. For such cues, we here introduce the term ‘*stress metabolites*’ as defining such compounds released, voluntarily or not, by an organism in response to abiotic stressors. These can be detected and processed by conspecifics and/or heterospecifics, leading to the induction of a behavioural stress response (Hazlett, 1985). Such a mechanism of indirect stress propagation might constitute positive feedback loops, defined as the propagation of stress responses within or between species by means of stress metabolites. Detecting stress metabolites may allow other individuals to modify their behaviour to avoid a change in the abiotic environment, and communities to coordinate or potentiate their response to ensure survival (Mothersill et al., 2007; Giacomini et al., 2015; Abreu et al., 2016). However, to investigate the function of any pH drop-induced chemical cues, the original stressor (pH drop) must first be removed from the experimental design.

In this study, we investigate the indirect effects of pH drop through stress metabolites induced by it on fitness-relevant behaviours within and among species inhabiting the intertidal zone. Using a full factorial design, we test the hypothesis that an acute pH drop to end-of-century level (7.6) will both directly and indirectly affect behaviour through the induction of stress metabolites. We observed three marine invertebrates (small hermit crab *Diogenes pugilator*, green shore crab *Carcinus maenas*, and harbour ragworm *Hediste diversicolor*) exposed to control pH and pH drop in combination with conditioned water from both conspecifics and their potential vertebrate predator, the gilt-head sea bream *S. aurata*.

## Methods

### Experimental design

Small hermit crabs *Diogenes pugilator* and green shore crabs *Carcinus maenas* were collected in autumn at low tide in the Ria Formosa lagoon (Portugal). Harbour ragworms (*Hediste diversicolor*) were sourced via a local supplier (Valbaits) from the Setubal lagoon (Portugal). Animals were acclimated for one week at pH 8.2 in large communal tanks mimicking natural habitats at the Ramalhete Marine Station (CCMAR, Faro, Portugal) in a direct CO_2_-controlled system with *p*CO_2_ constantly measured and adjusted as described in Sordo et al. (2016). Seawater parameters were measured daily (mean temperature: 18.90°C ± 0.97°C, mean salinity: 35.58 ± 0.19, mean dissolved oxygen: 91.60% ± 2.15%). A total pool of approx. 100-120 *D. pugilator*, 150-200 *C. maenas*, and over 150 *H. diversicolor* was used during the experimental period, yielding a total of 921 observations. Because no individual marking could be achieved, we aimed to space out the reuse of animals in successive behavioural assays by circulating animals through recovery tanks afterwards, before returning them to their stock population tanks once a day. Animal reuse was randomized across treatments and treatments randomized per day to prevent any confounding effects on the measured behaviours. All experiments were conducted in October 2019, except one independent set of observations (n = 40) in *C. maenas* conducted in September 2018, which was analysed together with the 2019 data with the year treated as a covariate.

A three-way factorial design of pH drop x stress metabolites x donor species was generated to study the effects of acute pH drop, stress metabolites induced by it, and their combination versus a control containing regular metabolites (control metabolites) (Figure 1). Metabolite donor species were *D. pugilator*, *C. maenas*, *H. diversicolor* (called conspecific donors) and the potential predator *Sparus aurata* (called heterospecific donor). Metabolite donors were conditioned for 30 minutes in seawater at regular pH (pH 8.2, 400 μatm CO_2_, putatively inducing control metabolite release), or pH drop (pH 7.6, 700 μatm CO_2_, putatively inducing stress metabolite release). Recipient species were *D. pugilator, C. maenas,* and *H. diversicolor* which received water containing either conspecific or heterospecific metabolites). To achieve the full factorial design, recipient species received either control or stress metabolites and were tested in either control pH (8.2) or pH drop (pH = 7.6), by re-adjusting the pH of the conditioned water before each bioassay. Overall, four different experimental treatments were obtained: (i) putative control metabolites at pH 8.2 (CM), (ii) putative control metabolites at pH 7.6 (pH drop), (iii) putative stress metabolites at pH 8.2 (SM), (iv) and putative stress metabolites at pH 7.6 (pH drop+SM). Testing conspecific and heterospecific metabolites with three recipient species amounted to a total of twelve treatments. Prior to use in recipient species, conditioned seawater from *H. diversicolor* and *S. aurata* (0.0067 fish/L) was tenfold diluted in fresh system seawater at the desired pH. To explore the possibility that stress metabolites are equivalent to highly concentrated control metabolites, we additionally tested the effects of undiluted *S. aurata* control metabolites at pH 8.2 in *H. diversicolor* and *D. pugilator*. See supplementary information for further details on treatments (Table S1) and water conditioning (supplementary methods).

**Figure 1.**
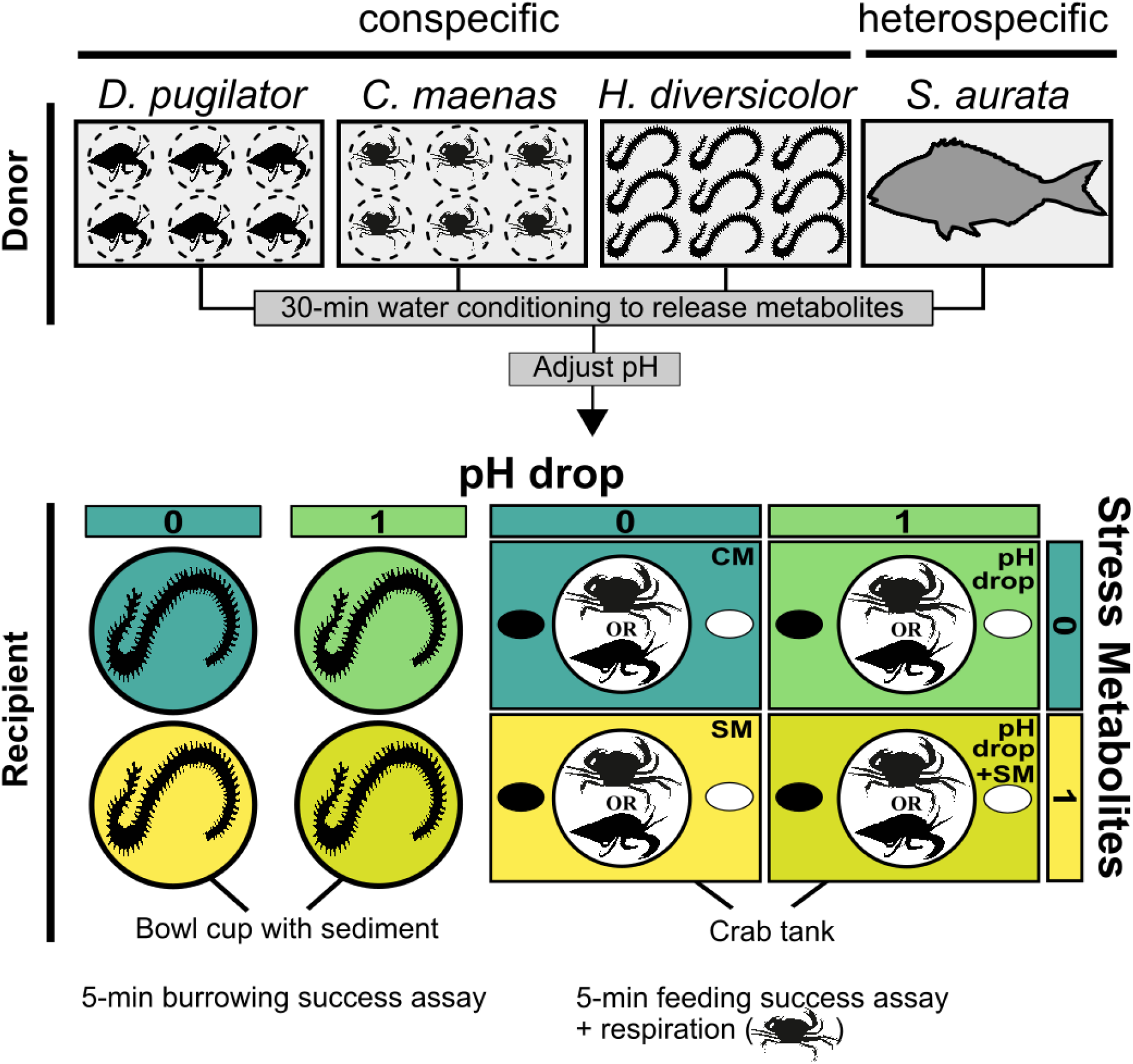
Schematic experimental design showing the presence vs. absence of predictors (pH drop, stress metabolites, donor). Behavioural effects of short-term acute pH drop and stress metabolites it induced were tested in three species: *Diogenes pugilator*, *Carcinus maenas*, and *Hediste diversicolor*. Metabolites were obtained by conditioning donors in either control pH = 8.2 (releasing putative control metabolites), or pH drop = 7.6 (releasing putative stress metabolites), followed by pH adjustments for factorial design yielding four experimental conditions CM, pH drop, SM, and pH drop+SM. Predictors are binary coded as 0 (control metabolites, control pH) and 1 (stress metabolites, pH drop). Metabolites originated from conspecifics or the heterospecific *Sparus aurata*. Behaviour assays for crabs consisted in locating a feeding cue in 300 seconds. Specimens of *H. diversicolor* were placed on top of sediment and burrowing behaviour was recorded for 300 seconds. A range of avoidance behaviours were also recorded. After completing the feeding assay, *C. maenas* additionally underwent five minutes of respiration rate measurements. Experimental conditions were CM: control metabolites in control pH, SM: control metabolites in control pH, pH drop: control metabolites in pH drop, pH drop+SM: stress metabolites in pH drop. Animal drawings by A. Murcia and KCWV.

### Behavioural bioassays and respirometry

For each of the four conditions, we measured the time to find a feeding cue (1/10 diluted mussel juice) *vs.* a mock (seawater) cue in *D. pugilator* and *C. maenas*, or to bury the head entirely in the sand in *H. diversicolor*, and dubbed this variable ‘time-to-success’. Behavioural assays were terminated once the animal successfully grabbed the ballasted sponge containing the feeding cue with their pincers (*D. pugilator* and *C. maenas*) or buried its entire head (*H. diversicolor*), or at a maximum time of 300 seconds. Both feeding and burrowing behaviours are tested response variables in crabs (de la Haye et al., 2012; Wang et al., 2018) and ragworm (Bhuiyan et al., 2021) exposed to pH drop. Additionally, avoidance behaviour was binary coded. In *D. pugilator* and *C. maenas*, these avoidance behaviours included freezing (suddenly retreating into the shell or attempting to burrow, sudden and lasting arrest in locomotion), and escaping (walking along the walls or retreating into a corner of the tank). Such freeze and escape behaviours indicate danger in crustaceans (Katz and Rittschof, 1993; Perrot-Minnot et al., 2017; Tomsic et al., 2017). In *H. diversicolor*, avoidance behaviours consisted of freezing, which might be accompanied by spread of jaws, sideways-undulating behaviour, and formation of a noticeable slime cap whilst outside the sediment. These behaviours indicate extreme stress in marine polychaetes (Mouneyrac et al., 2003; Burlinson and Lawrence, 2007; McBriarty et al., 2018). The feeding cues were randomly assigned left or right to the crabs. Additionally, respiration rates of *C. maenas* were monitored immediately after their use in feeding behaviour assays in a two-way factorial design of pH drop x stress metabolites using only conspecific donor metabolites. See supplementary methods for further details on behavioural assays and respirometry.

### Statistical analysis

Avoidance behaviours were analysed using generalised linear models for logit regression with binomial distribution using the *stats* R package (R Core Team, 2020). Time-to-success was modelled for the time to reach a feeding cue in *D. pugilator* and *C. maenas* or to burrow the head in *H. diversicolor* depending on the three predictors: pH drop, stress metabolites, and donor. Time-to-success data was analysed with a time-to-event analysis (also called survival analysis) using Cox proportional hazard models from the *survival* R package (Therneau and Grambsch, 2000; Therneau, 2021). The event being the success of food cue location (in crabs) or burrowing the head in sediment (in worms), the analysis was hereafter referred to as ‘time-to-success analysis’. The exponentiated estimates (hazard ratios) from the Cox proportional hazard models were expressed as ‘success ratio’. Success ratios were visualised using the *plot_model* function from the *sjPlot* R package (Lüdecke, 2021). The ‘success probability’ (the probability of the feeding or burrowing event occurring at any time over 300s) over time was represented by Kaplan-Meier curves drawn using the *survminer* R package (Kassambara et al., 2020). Animals not reaching the feeding cue or not burying their head were censored and assigned the maximum time of ‘300+’. The oxygen measurements were transformed using the additive-inverse slope coefficients standardised to the carapace width and rescaled to the mean additive-inverse slope coefficient of the control CM as shown in equation (1) to yield respiration rates.

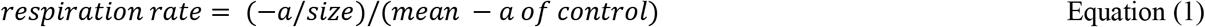

where a is the slope of the oxygen levels (%) as a function of time (s) and the size is proxied by the carapace width.

All statistical models always included the main effects and interaction terms of the binary predictors pH drop (pH = 8.2 vs. pH = 7.6), stress metabolites (stress metabolites vs. control metabolites), and donor (conspecific vs. heterospecific). Covariates (year, number of water uses, crab size, where relevant) were included in models when deemed significant compared to the null model by an ANOVA Chi-square test from the *stats* R package (R Core Team, 2020). Overall model fit p-values were retrieved from the ANOVA Chi-squared test or the Likelihood ratio test for generalised linear models and Cox-proportional models, respectively. Pairwise comparisons between treatments involved in the interaction term between the three predictors were obtained from statistical models using the *emmeans* R package (Lenth, 2021), wherein false discovery rate p-value adjustments were applied for post-hoc term-wide multiple testing.

To investigate the possibility that stress metabolites are equivalent to highly concentrated control metabolites (which were present in CM and pH drop treatments but in tenfold dilution), additional tests compared the behavioural response of *D. pugilator* and *H. diversicolor* to tenfold diluted stress metabolites and control metabolites to that of undiluted control metabolites from *S. aurata*.

Effect sizes of the respiration rates were measured as Cohen’s d using the effsize R package (Torchiano, 2020) for estimates between two groups, or according to Lenhard and Lenhard (2016) for estimates of the interaction term between pH drop and stress metabolites predictors. Effect sizes were classified following the classification given in Sawilowsky (2009). All statistical analyses were conducted in RStudio (RStudio Team, 2020) with a significance threshold of P ≤ 0.05. See R script, datasets, and supplementary information for further details.

## Results

### *Response of small hermit crab* Diogenes pugilator *to pH, stress metabolite, and donor predictors*

The time-to-success analysis was not conclusive for the stress metabolite predictor in *D. pugilator* (Figure 2A, Table 1, Z = −0.66, P = 0.5073). On the other hand, the Cox proportional hazard model (overall Likelihood ratio test model fit: P = 0.06) found that both pH drop (feeding success ratio = 0.47, Z = −2.14, P = 0.0325) and metabolites from *S. aurata* (feeding success ratio = 0.35, Z = −2.76, P = 0.0058) had significant negative effects on the feeding success ratio of *D. pugilator.* In addition, there was a significant interaction between pH drop and metabolite donor terms on the feeding success of *D. pugilator* (Z = 2.19, P = 0.0289). Overall, pH drop induced a significantly lower success (risk) score in the conspecific treatment (Figure 2B). Post-hoc analyses evidenced that all treatments (CM, pH drop, SM, and pH drop+SM) induced similar feeding time responses in both the conspecific and heterospecific donor groups (Table S2).

**Figure 2.**
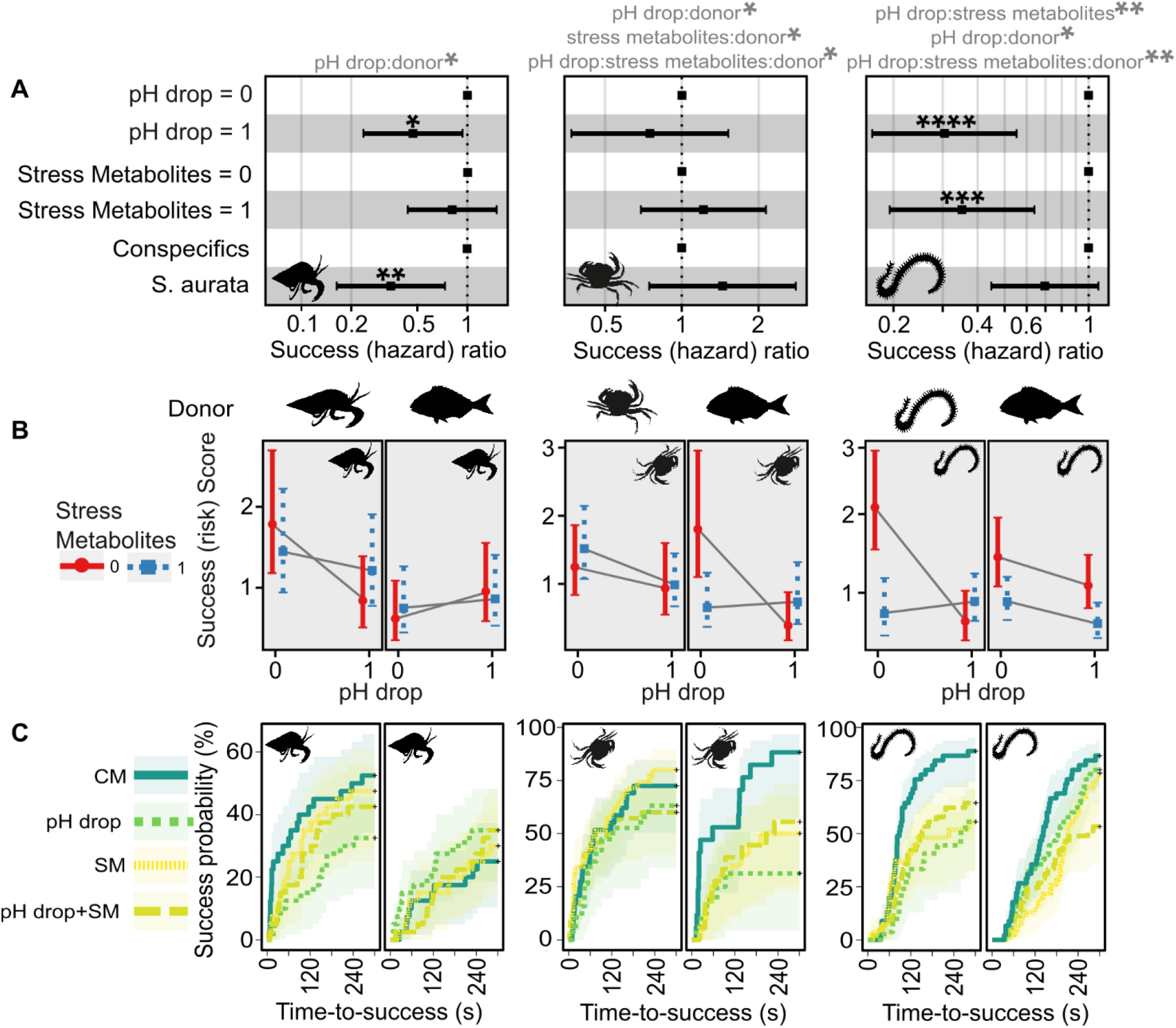
Effects of predictors (pH drop, stress metabolites, donor) on time-to-success in *Diogenes pugilator*, *Carcinus maenas*, and *Hediste diversicolor*. A) effects of predictors on success ratio (aka hazard ratio with success as event, arbitrary units). Likelihood ratio tests for overall model fit were: *D. pugilator*: P = 0.06; *C. maenas*: P = 0.009; *H. diversicolor*: P < 0.0001). Significant predictors from Cox proportional hazard models are shown with asterisks, and significant interaction terms are shown in grey above the plots. B) Interaction term plot of marginal effects showing predicted success score (aka risk score with success as event, arbitrary units ± confidence interval) split by donor and recipient species. Crossed solid grey lines represent an interacting effect of pH and metabolites. C) Kaplan-Meier curves to visualise success probability (cumulative event) for each experimental condition over time. *: P ≤ 0.05, **: P < 0.01, ***: P < 0.001, ****: P < 0.0001. Experimental treatments were: CM: control metabolites in control pH, pH drop: control metabolites in pH drop, SM: stress metabolites in control pH, pH drop+SM: stress metabolites in pH drop.

**Table 1.** Results of Cox proportional hazard model for the main effects of predictors (pH drop, stress metabolites, donor) on the time-to-success analysis in small hermit crab (*Diogenes pugilator*, n = 320 observations with 120 events of finding a food cue), green shore crab (*Carcinus maenas*, n = 189 observations with 122 events of finding a food cue), and harbour ragworm (*Hediste diversicolor*, n = 325 observations with 234 events of burrowing head in sediment). Unsuccessful observations are censored. Significance (P ≤ 0.05) is shown by p-values in bold. Overall significance of the models using Likelihood ratio tests were: *D. pugilator*: P = 0.06; *C. maenas*: P = 0.009; *H. diversicolor*: P < 0.0001). Covariates were dropped from models after analyses of deviance showed that they passed the Chi-squared test (*D. pugilator*: number of water uses: P = 0.1163, crab size: P = 0.6318; *C. maenas*: number of water uses: P = 0.6104, crab size: P = 0.9995, year: P = 0.3348; *H. diversicolor*: number of water uses: P = 0.2649). Success ratio (aka hazard ratio) is the exponentiated estimate. SE is the standard error of estimate.

| Predictors | Success (hazard) ratio | Estimate | SE | Z | P |
| --- | --- | --- | --- | --- | --- |
| *D. pugilator* |  |  |  |  |  |
| pH drop | **0.4702** | -0.7546 | 0.3530 | -2.1378 | **0.0325** |
| stress metabolites | 0.8106 | -0.2100 | 0.3167 | -0.6631 | 0.5073 |
| donor | **0.3460** | -1.0613 | 0.3843 | -2.7613 | **0.0058** |
| pH drop:stress metabolites | 1.7878 | 0.5810 | 0.4858 | 1.1959 | 0.2317 |
| pH drop:donor | **3.2842** | 1.1891 | 0.5441 | 2.1853 | **0.0289** |
| stress metabolites:donor | 1.4970 | 0.4034 | 0.5326 | 0.7575 | 0.4487 |
| pH drop:stress metabolites:donor | 0.4168 | -0.8752 | 0.7498 | -1.1672 | 0.2431 |
| *C. maenas* |  |  |  |  |  |
| pH drop | 0.7490 | -0.2890 | 0.3620 | -0.7982 | 0.4247 |
| stress metabolites | 1.2156 | 0.1953 | 0.2887 | 0.6762 | 0.4989 |
| donor | 1.4445 | 0.3678 | 0.3385 | 1.0865 | 0.2773 |
| pH drop:stress metabolite | 0.8674 | -0.1422 | 0.4630 | -0.3072 | 0.7587 |
| pH drop:donor | **0.2870** | -1.2484 | 0.6312 | -1.9777 | **0.0480** |
| stress metabolites:donor | **0.2981** | -1.2102 | 0.5008 | -2.4165 | **0.0157** |
| pH drop:stress metabolites:donor | **6.0273** | 1.7963 | 0.8261 | 2.1743 | **0.0297** |
| *H. diversicolor* |  |  |  |  |  |
| pH drop | **0.3043** | -1.1899 | 0.3045 | -3.9075 | **< 0.0001** |
| stress metabolites | **0.3520** | -1.0442 | 0.3049 | -3.4251 | **0.0006** |
| donor | 0.6965 | -0.3616 | 0.2258 | -1.6013 | 0.1093 |
| pH drop:stress metabolites | **3.9914** | 1.3841 | 0.4411 | 3.1376 | **0.0017** |
| pH drop:donor | **2.4669** | 0.9030 | 0.3816 | 2.3663 | **0.0180** |
| stress metabolites:donor | 1.7318 | 0.5492 | 0.3816 | 1.4390 | 0.1501 |
| pH drop:stress metabolites:donor | **0.2291** | -1.4736 | 0.5633 | -2.6158 | **0.0089** |

Avoidance responses of *D. pugilator* did not depend on pH drop (Z = 0.25, P = 0.8049) nor stress metabolites (Z = −0.25, P = 0.7995, overall Chi-square test model fit: P < 0.0001, Table 2), but were significantly more pronounced when metabolites originated from *S. aurata* (69%) instead of conspecifics (26%, Z = 2.46, P = 0.0139, Figure 3, Table 2). After splitting the display of avoidance behaviour by donor, pairwise comparisons failed to find differences between the four treatments (CM, pH drop, SM, and pH drop+SM), but confirmed that the donor effect existed across all four treatments (Table S3, Figure 3). However, partitioning the avoidance behaviour of *D. pugilator* into freezing and escaping responses evidenced that escaping significantly increased with *S. aurata* control metabolites only when tested in pH drop (Figures S1-S2, Tables S4-S7, see supplementary results).

**Table 2.** Results of the binomial generalised linear model for the main effects of predictors (pH drop, stress metabolites, donor) on the avoidance behaviour in small hermit crab (*Diogenes pugilator*, n = 320 observations of finding a food cue), green shore crab (*Carcinus maenas*, n = 151 observations of finding a food cue), and harbour ragworm (*Hediste diversicolor*, n = 325 observations of burrowing head in sediment). Due to missing observations in conspecific control metabolites at control pH in Hediste diversicolor, data was analysed with two models. Model 1: effect of donor across all treatments (sample size: 253 observations). Model 2: effects of pH and metabolites in subset receiving *S. aurata* metabolites (sample size: 154 observations). Overall significance of models from Chi-squared analyses of deviance when including only predictors were: *D. pugilator*: P < 0.0001; *C. maenas*: P < 0.0001; *H. diversicolor*: P < 0.0001 (model 1) and P < 0.0001 (model 2). Covariates were dropped from models after analyses of deviance showed that they passed the Chi-squared test (*D. pugilator*: number of water uses: P = 0.5963, crab size: P = 0.9158; *C. maenas*: crab size: P = 0.2710). Significance (P ≤ 0.05) is shown by p-values in bold. SE is the standard error of estimate.

| Predictors | Estimate | SE | Z | P |
| --- | --- | --- | --- | --- |
| ***D. pugilator*** |  |  |  |  |
| (Intercept) | -0.9694 | 0.3541 | -2.7380 | 0.0062 |
| pH drop | 0.1221 | 0.4944 | 0.2470 | 0.8049 |
| stress metabolites | -0.1292 | 0.5086 | -0.2540 | 0.7995 |
| donor | 1.1701 | 0.4758 | **2.4590** | **0.0139** |
| pH drop:stress metabolites | -0.2603 | 0.7219 | -0.3610 | 0.7185 |
| pH drop:donor | 0.7758 | 0.6919 | 1.1210 | 0.2622 |
| stress metabolites:donor | 1.0272 | 0.7022 | 1.4630 | 0.1435 |
| pH drop:stress metabolites:donor | -0.8890 | 1.0039 | -0.8860 | 0.3759 |
| ***C. maenas*** |  |  |  |  |
| (Intercept) | -0.2224 | 0.6725 | -0.3307 | 0.7409 |
| pH drop | -1.1263 | 0.9173 | -1.2279 | 0.2195 |
| stress metabolites | -1.1263 | 0.9173 | -1.2279 | 0.2195 |
| donor | -0.4626 | 0.8326 | -0.5556 | 0.5785 |
| number of water uses | -0.3077 | 0.1556 | **-1.9772** | **0.0480** |
| pH drop:stress metabolites | 2.2527 | 1.2978 | 1.7357 | 0.0826 |
| pH drop:donor | 5.5551 | 1.5392 | **3.6090** | **0.0003** |
| stress metabolites:donor | 3.8799 | 1.2465 | **3.1127** | **0.0019** |
| pH drop:stress metabolites:donor | -7.5415 | 1.9329 | **-3.9017** | **0.0001** |
| ***H. diversicolor*** |  |  |  |  |
| Model 1 (both donors) |  |  |  |  |
| (Intercept) | 0.2801 | 0.3790 | 0.7391 | 0.4598 |
| donor | 1.0944 | 0.3169 | **3.4534** | **0.0006** |
| number of water uses | -0.7745 | 0.1462 | **-5.2980** | **< 0.0001** |
| Model 2 (*S. aurata* donor) |  |  |  |  |
| (Intercept) | -0.6931 | 0.5000 | -1.3863 | 0.1657 |
| pH drop | -18.8729 | 1603.1137 | -0.0118 | 0.9906 |
| stress metabolites | 3.3557 | 0.7788 | **4.3086** | **< 0.0001** |
| pH drop:stress metabolites | 16.2548 | 1603.1138 | 0.0101 | 0.9919 |

**Figure 3.**
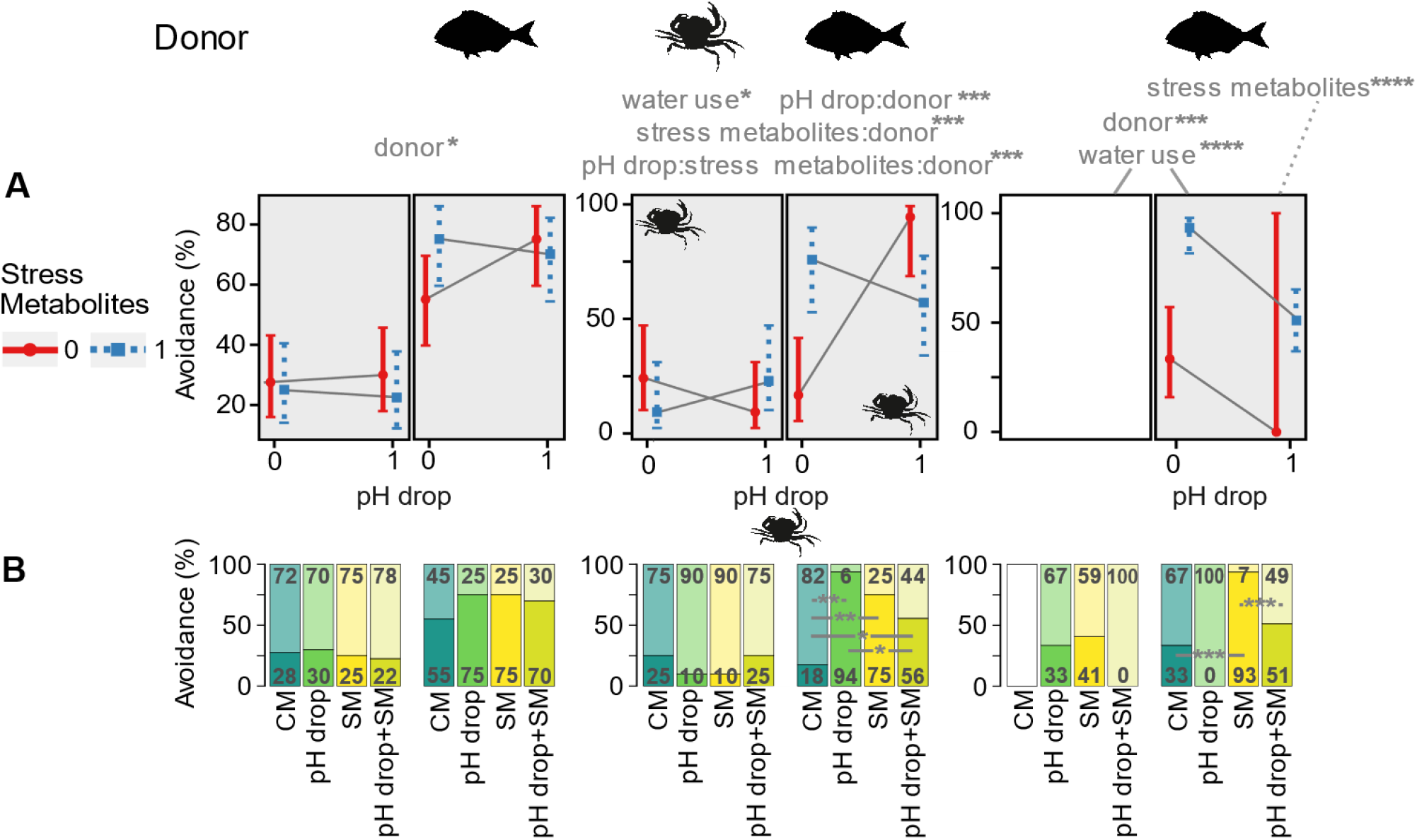
Effects of predictors (pH drop, stress metabolites, donor) on the percentage of avoidance behaviour in *Diogenes pugilator, Carcinus maenas, and Hediste diversicolor*. Avoidance behaviour included freezing and escaping (*D. pugilator* and *C. maenas*), or freezing, curling, flipping, and slime secretion (*H. diversicolor*). A) Split bars represent the presence (dark area) or absence (light area) of avoidance behaviours. Significant main predictors (and their interaction terms) and covariates are shown above plots. Significant pairwise comparisons between treatments in each donor/recipient are shown as horizontal grey lines. *: P ≤ 0.05, **: P < 0.01, ***: P < 0.001, ****: P < 0.0001. Experimental treatments were: CM: control metabolites in control pH, pH drop: control metabolites in pH drop, SM: stress metabolites in control pH, pH drop+SM: stress metabolites in pH drop. (B) Interaction plot showing the marginal effects on the predicted avoidance percentages (± confidence interval) of stress metabolites and pH drop within the conspecific and heterospecific groups for each species. Crossing solid grey lines represent an interacting effect of pH and metabolites for each donor. The interactive effects of predictors for *H. diversicolor/H. diversicolor* are not represented due to the missing CM treatment. Overall effects of donor and water use across the response of *H. diversicolor* (for either donor) are shown with solid lines, whereas the effects of stress metabolites are for *S. aurata/H. diversicolor* only (grey dashed line).

### *Response of green shore crab* Carcinus maenas *to pH, stress metabolite, and donor predictors*

The Cox proportional hazard model (overall Likelihood ratio test model fit: P = 0.009) showed that the feeding success ratio of *C. maenas* did not significantly vary with pH (Z = −0.80 P = 0.4247), metabolites (Z = 0.68, P = 0.4989), nor donor (Z = 1.09, P = 0.2773) terms (Figure 2, Table 1). Nevertheless, all interaction terms were significant. Post-hoc tests revealed that time-to-success responses were similar across treatments CM, pH drop, SM, and pH drop+SM when *C. maenas* received conspecific metabolites (Table S8). This pattern changed when *S. aurata* was the metabolite donor as evidenced by significantly lower feeding success scores in *C. maenas* exposed to pH drop (Z = 2.97, P = 0.0179), SM (Z = 2.48, P = 0.0394), and the trend for pH drop+SM (Z = 2.20, P = 0.0563), compared to the control CM.

In *C. maenas*, the predictors pH drop (Z = −1.23, P = 0.2195), stress metabolites (Z = −1.23, P = 0.2195), and donor (Z = −1.23, P = 0.5785) did not alter avoidance responses but their interaction terms were significant (overall Chi-square test model fit: P < 0.0001, Table 2, Figure 3). Pairwise comparisons showed no differences between the four treatments (CM, pH drop, SM, pH drop+SM) in avoidance patterns of *C. maenas* receiving conspecific metabolites. Conversely, *C. maenas* facing *S. aurata* metabolites while experiencing treatments of pH drop (94%, Z = −3.60, P = 0.0020), SM (75%, Z = −3.28, P = 0.0031), and pH drop+SM (56%, Z = −2.33, P = 0.04) significantly increased their avoidance display compared to control CM (18%). Moreover, pH drop combined with stress metabolites (pH drop+SM treatment) instead of control metabolites (pH drop treatment) significantly lowered avoidance behaviours (Z = 2.21 P = 0.0405, Table S9, Figure 3). Partitioning the avoidance behaviour of *C. maenas* into freezing and escaping responses evidenced that escaping explained the avoidance response whereas no clear patterns were observed in the freezing behaviour (Figures S1-S2, Tables S10-S13, see supplementary results). Lastly, neither pH drop nor stress metabolites altered the respiration rates of *C. maenas* although the interaction was near to significant (P = 0.0531, Figure S3, Table S14).

### *Response of harbour ragworm* Hediste diversicolor *to pH, stress metabolite, and donor predictors*

Burrowing success responses of *H. diversicolor* did not depend on the donor species (P = 0.1093, Figure 2, Table 1). However, the Cox proportional hazard model (overall Likelihood ratio test model fit: P < 0.0001) evidenced that both pH drop (burrowing success ratio = 0.30, Z = −3.91, P < 0.0001) and stress metabolites (burrowing success ratio = 0.35, Z = −3.42, P = 0.0006) terms significantly altered burrowing time responses of *H. diversicolor*. Significant interactions between the main predictors required to decipher the involvement of within-donor treatment effects (Table S15). Post-hoc tests revealed that pH drop (Z = 3.91, P = 0.0006), SM (Z = 3.43, P = 0.0012), and pH drop+SM (Z = 3.47, P = 0.0012) treatments significantly lowered burrowing success scores compared to the control CM in *H. diversicolor* exposed to conspecific metabolites. Moreover, *H. diversicolor* facing *S. aurata* metabolites had lower burrowing success scores in the pH drop+SM treatment compared to the control CM (Z = 3.35, P = 0.0049).

*H. diversicolor* displayed more avoidance in response to *S. aurata* metabolites (47%) compared to conspecific metabolites (20%, Z = 1.09, P = 0.0006, overall Chi-square test model fit: P < 0.0001, Table 2, Figures 3). *H. diversicolor* exposed to *S. aurata* metabolites were not affected by pH drop (Z = −0.01, P = 0.9906) whereas stress metabolites induced significantly more avoidance than control metabolites (Z = −5.30, P < 0.0001, overall Chi-square test model fit: P < 0.0001, Table 2). There were no differences across pH drop, SM, and pH drop+SM treatments in *H. diversicolor* receiving conspecific metabolites (Table S16). Pairwise comparisons revealed that the SM treatment (93%) triggered significantly more avoidance than CM (33%, Z = −4.31, P = 0.0001) and pH drop+SM treatments (51%, Z = 3.92, P = 0.0003). Specimens of *H. diversicolor* exposed to *S. aurata* metabolites displayed different avoidance responses in the control treatment CM (video 1) compared to pH drop (slowed-down burrowing response, video 2), SM (sideways-undulating behaviour with body flips, video 3), and pH drop+SM (raised head and spread mouth parts with freezing behaviour in video 4 and slowed-down movement with sideways undulating behaviour in video 5).

## Discussion

### Direct effects of pH drop and donor type on assay success and avoidance behaviours

Changes in fitness-relevant behaviours to acute pH change can be expected to show high variance and small effect sizes (Clark et al., 2020a), and may therefore require large observational datasets. In this study, we performed over 900 behavioural observations, and our results reflect this high variance. *Sparus aurata-* conditioned water, whether it contained control metabolites or pH stress-induced metabolites, elicited avoidance in all recipient species which can be attributed to antipredator behaviours. Despite the donor effect however, much of the behavioural variance was explained by the effects of pH drop and stress metabolites, their interaction, or their interaction with the donor term (Figure 2).

We found that pH drop lowered success ratios in feeding and burrowing in *Diogenes pugilator* and *Hediste diversicolor*, respectively. Such low pH-induced impairments of fitness-relevant behaviours were previously found in hermit crabs *Pagurus bernhardus* and *P. tanneri* and may indicate impaired decision-making (de la Haye et al., 2011, 2012; Kim et al., 2015). Past literature also reported delayed burrowing responses in the harbour ragworm *H. diversicolor* exposed for 28 days to pH drop (Bhuiyan et al., 2021), and an inverse relationship between burrowing and pH in the king ragworm *Alitta virens* (Batten and Bamber, 1996). These behavioural alterations may reflect the cost of acid-base regulation to counteract hypercapnia (Pörtner, 2008), as marine ragworms overexpress the acid-base regulator *Carbonic Anhydrase* (*CA*) at the expense of energy reserves in response to acidification (Freitas et al., 2016; Wage et al., 2016). In the longer term, behavioural costs of an altered burrowing activity may extend to crucial ecological functions as shown by the diminished bioturbation activity of *H. diversicolor* in similar conditions (Bond, 2018). In contrast, *Carcinus maenas* did not delay its feeding response to pH drop in presence of conspecific metabolites, which supports recent observations of ocean acidification having weaker effects than previously expected on marine animal behaviour (Clark et al., 2020a; Clements et al., 2020a, 2020b, 2021) including feeding responses in aquatic arthropods (Clements and Darrow, 2018). The resilience to pH stress in *C. maenas* may be due to their extracellular pH regulatory capacities allowing them to maintain feeding during short-term assays (Appelhans et al., 2012). Therefore, short-term hypercapnia is not a strong stressor in *C. maenas*, due to its pH-variable intertidal habitat (Fehsenfeld et al., 2011), which is supported by its ability to maintain respiration rates in our study. Other than reduced success ratios, pH drop itself did not induce any avoidance behaviours indicative of stress.

Besides the direct effect of pH drop on behaviour, previous studies on ocean acidification also found contrasting effects of pH drop on predator-prey interactions (Draper and Weissburg, 2019). Here, we observed an increased escaping tendency of *C. maenas* in response to *S. aurata* control metabolites in pH drop but not in control pH. Although this was not the main focus of our study, it could mean that pH drop renders predator odour more potent for its prey, for example through pH-dependent changes in odour molecular structure, receptor binding, or information processing (Munday et al., 2009; Roggatz et al., 2016, 2019; Schirrmacher et al., 2020).

### Indirect effects of pH drop on recipients’ behaviour through stress chemical communication

Despite the above-mentioned species-specific direct effects of pH drop on behaviour success ratios, we could show here that pH drop also has indirect effects on behaviour by altering chemical communication. We found that stress metabolites released by animals exposed to pH drop altered feeding and burrowing, and increased avoidance behaviours, but that the response was donor and recipient species-specific. Hermit crabs *D. pugilator* did not react to any stress metabolites. Significant responses in the other two recipient species, however, were directionally equivalent to those induced by pH drop itself since it took longer to find a food cue or to burrow the head in the presence of stress metabolites. Although behavioural stress response propagation through chemical communication is well documented for alarm substances and disturbance cues upon predation stress (Giacomini et al., 2015; Abreu et al., 2016; Mathuru, 2016), our results show here that chemical communication induced by abiotic stressors such as pH drop can trigger the release of waterborne chemicals that in turn trigger a behavioural stress response in its recipients. Stress metabolites conditioned by the potential predator *S. aurata* induced more avoidance behaviours and delayed feeding in *C. maenas* and burrowing in *H. diversicolor,* compared to *S. aurata* control metabolites. Similarly, the presence of conspecific stress metabolites significantly lowered *H. diversicolor* burrowing success ratio compared to conspecific control metabolites.

On the other hand, *C. maenas* did not react to conspecific stress metabolites. The different behavioural effect of conspecific stress metabolites between crabs and *H. diversicolor* has several possible explanations ranging from different defence strategies related to stress risk perception and evaluation (Hazlett, 1985; Bairos-Novak et al., 2017, 2018; Goldman et al., 2020b) to different types, concentrations, or ratios of the released chemicals (Júnior et al., 2010; Morishita and Barreto, 2011). As we have shown here a pH drop to 7.6 is not a strong stressor in crabs (Fehsenfeld et al., 2011) which may explain that we failed to observe responses to water from pH drop-conditioned conspecific green shore crabs. However, the presence of stress metabolites conditioned by *S. aurata* caused a marked drop in the success ratio of *C. maenas* at low (but not control) pH, evidenced by significant interaction terms.

Whilst we did not investigate their chemical structure, our experiment allows us to characterize novel properties of stress metabolites. Stress metabolites, like disturbance cues, potentially consist of regularly excreted metabolites such as urea and ammonia (Bairos-Novak et al., 2017; Shrivastava et al., 2019). The fact that behavioural responses of *H. diversicolor* and *D. pugilator* remained unchanged even when we used undiluted control metabolites (Tables S17-S18), might mean that stress metabolites are not just up-concentrated control metabolites. We also observed that avoidance behaviours by both *C. maenas* and *H. diversicolor* became less pronounced with subsequent water uses, showing that the cues are either volatile or have a very short half-life (less than half a day) in seawater. Alternatively, recipients may be able to discriminate degraded versus fresh metabolites and react accordingly to the degree of threat they may indicate (Fuselier et al., 2009; Bairos-Novak et al., 2018). This shows that stress metabolites induced by pH drop are either not identical with metabolites released in normal pH, and/or are at least more than tenfold higher concentrated than control metabolites.

The avoidance behaviours we recorded in this study are known indicators of stress. *S. aurata* stress metabolites elicited atypical freezing, eversion of the proboscis, mucus secretion, flipping and sideways-undulating behaviours in *H. diversicolor* which were previously described as indicators for a physiological stress response following exposure to copper sulphate (Burlinson and Lawrence, 2007), antiparasitic drugs (McBriarty et al., 2018), and trace metals (Mouneyrac et al., 2003). We also observed increased freeze and escape responses in crabs, which indicate a stress response in crustaceans (Katz and Rittschof, 1993; Perrot-Minnot et al., 2017; Tomsic et al., 2017).

The finding that *S. aurata* pH-induced stress metabolites induced such responses in potential prey species suggests that responses to disturbance cues are not depending on the ‘audience’, similar as what has been observed in tadpoles (Bairos-Novak et al., 2020). In our study, stress metabolites from *H. diversicolor* altered the burrowing success of conspecifics whereas Watson and colleagues (2005) found that whole-body extracts of *H. diversicolor* did not alter out-of-burrow activities of the king ragworm *Alitta virens* (of the same subfamily Nereidinae), indicating that stress metabolites may also differ from alarm cues.

A directionally similar response to pH drop and to the stress metabolites it induces shows that pH drop can have indirect effects, reminiscent of a positive feedback loop, by which stressed animals negatively influence fitness-relevant behaviours of community members. This was first observed by Hazlett (1985) after freshwater crayfish exposed to water conditioned with heat-stressed conspecifics displayed increased alertness, after returning the conditioned water to normal temperature. Similar indirect effects through chemical signalling from stressed donors to unstressed recipients were also shown in response to different types of biotic stressors such as mock predator chase (Toa et al., 2004; Giacomini et al., 2015), handling (Barcellos et al., 2011), and acute fasting (Abreu et al., 2016), and to physical injury including irradiation (Mothersill et al., 2007) and predation (Frisch, 1938; Mathuru et al., 2012; Oliveira et al., 2013).

## Conclusions and Perspectives

In this study, we could show that short-term pH drop of a similar magnitude of that experienced within the intertidal zone, but also aligned to end-of-century predicted average values (Chavez et al., 2017; Landschützer et al., 2018), had negative consequences on fitness-relevant behaviours in harbour ragworm *H. diversicolor* and small hermit crab *D. pugilator*. Additionally, we confirm that pH drop events also impede the same behaviours in the same way indirectly via chemical communication, albeit these effects depended on donor and recipient species. Sea bream *S. aurata* and harbour ragworm *H. diversicolor* stressed by pH drop released stress metabolites which likely differ from control metabolites and negatively affect fitness-relevant behaviours in metabolite recipients. Stress metabolites induced similar avoidance behaviours as those exhibited under physiological stress, meaning that a stress response was propagated from donor to recipient. These negative indirect effects, or positive feedback loops, warrant further study, especially as our results were inconclusive with regards to the combined treatments of low pH and stress metabolites. Short-term pH drops thus involve behavioural additionally to metabolic trade-offs, which is also of interest for better predicting the response of natural and aquaculture systems under ocean acidification combined with tidal pH fluctuations. We are hopeful that our more than 900 observations and balanced experimental design could overcome most potential limitations, in relation to pseudoreplication due to reusing some animals on different days, and any overestimation of effect sizes related to small sample size (Clements et al., 2020b). Given the potential high ecological significance of indirect negative effects of pH fluctuations on population fitness, studies aiming to replicate our results should aim to further optimise our experimental design with better technical equipment as that available to us (Baker, 2016).

## Ethics

All experiments were approved by the University of Hull Ethics Committee under the approvals U020 and FEC_2019_81.

## Authors’ contributions

KWV designed the study. Experiments were performed by LF, SS, LA, JF, RN, KR, and KWV. LF analysed the data and wrote the first manuscript draft with KWV, JDH and HBH. All authors contributed to the final manuscript.

## Acknowledgements

The authors wish to acknowledge J. Pena dos Reis, P. Hubbard, Z. Velez, and all staff members of the CCMAR-Ramalhete Marine Station. JDH and KWV acknowledge funding by Natural Environment Research Council (NERC grant #NE/T001577/1), and KWV, JDH and LF acknowledge funding by the University of Hull towards the MolStressH2O research cluster. We thank Andrea Murcia for the drawings of *Diogenes pugilator*, *Sparus aurata*, and *Hediste diversicolor*.

