## supplementary information for "Behavioural stress feedback loops in benthic invertebrates caused by pH drop-induced metabolites"

**Supplementary Material 1 containing the Supplementary Information from 'Behavioural stress feedback loops in benthic invertebrates caused by pH drop-induced metabolites'.**

Lauric Feugere[1], Lauren Angell[1], James Fagents[1], Rebecca Nightingale[1], Kirsty Rowland[1], Saffiyah Skinner[1], Jorg Hardege[1], Helga Bartels-Hardege[1], Katharina C. Wollenberg Valero[1]*

[1] Department of Biological and Marine Sciences, University of Hull, Cottingham Road, Kingston-upon-Hull HU67RX, United Kingdom.

### Supplementary Methods

Details of water conditioning with metabolites

To condition water for experiments, each nine harbour ragworms *Hediste diversicolor*, six small hermit crabs *Diogenes pugilator*, six green shore crabs *Carcinus maenas*, or one gilt-head sea bream *Sparus aurata* (~ 880 g) were placed in separate tanks with 0.015 L, 1 L, 5 L, or 15 L of seawater, respectively. Water conditioned by *S. aurata* was provided by the Ramalhete Marine Station. Specimens of *D. pugilator* and *C. maenas* were separated into transparent mesh-bottomed plastic cups to prevent intraspecific interactions while allowing any metabolites to diffuse into the seawater. Specimens of *H. diversicolor* were placed together on top of the sediment into which they tended to burrow individually. Metabolite donor and recipient species couples in the conspecific groups were *D. pugilator/D. pugilator*, *C. maenas/C. maenas*, and *H. diversicolor/H. diversicolor*. Metabolite donor and recipient species couples in the heterospecific groups were: *S. aurata/D. pugilator*, *S. aurata/C. maenas*, and *S. aurata/H. diversicolor*. Conditioning of water with metabolites took place for 30 min at either regular pH (pH 8.2, 400 µatm $CO_2$, putatively inducing control metabolite release), or pH drop (pH 7.6, 700 µatm $CO_2$, putatively inducing stress metabolite release). Conditioned water containing metabolites from *H. diversicolor* and *S. aurata* was tenfold diluted prior use in recipient species. To achieve the full factorial design, control metabolites were tested in pH drop and stress metabolites were tested at control pH. For this purpose, the pH of control metabolite conditioned seawater was reduced to pH = 7.6 (using 0.1 M HCl for control metabolites conditioned by crabs, or by dilution in system water at pH = 7.6 for control metabolites conditioned by *H. diversicolor* and *S. aurata*). On the other hand, the pH of stress metabolite conditioned water was increased to pH = 8.2 (using reef buffer™ – Seachem, Georgia, USA – for control metabolites conditioned by crabs, or by dilution in system water at pH = 8.2 for control metabolites conditioned by *H. diversicolor* and *S. aurata*). Addition of reef buffer™ did not alter the salinity (measured with a salinity meter). The conditioning of the water is summarised in Table S1.

Details of behavioural experiments

Our preliminary experiments (in Autumn 2018, n = 40) showed no effect of the number of conditioned water uses until more than five uses on the time-to-success of *C. maenas* ($P = 0.373$), as well as a disinterest of male *C. maenas* in food cues in the mating period. We therefore limited experiments in *C. maenas* to females, and each batch of conditioned water was used for at most five behaviour assays before being renewed. Between each batch of conditioned water, sand in the bioassay tanks was thoroughly rinsed with system water.

The behaviour of *H. diversicolor* was tested using circular plastic bowl cups (H: 5 cm x ⌀: 9 cm) filled with 1 cm cleaned fine sand from the Ria Formosa lagoon and 15 mL conditioned seawater resulting in approx. 1 cm seawater layer above the sand. The behaviour of both *D. pugilator* and *C. maenas* was tested in binary choice sport-pitch-like tanks. Tanks used for *D. pugilator* (H: 17 cm x W: 15 cm x L: 25 cm) were filled either with 1 L conditioned seawater and a thin (~ 1 cm) layer of clean fine sand, as *D. pugilator* showed reluctance moving over plastic surfaces without substrate. Test tanks used for *C. maenas* (H: 24 cm x W: 19 cm x L: 37 cm) were filled with 5 L conditioned seawater and were free of sand. Both *D. pugilator* and *C. maenas* tanks were wrapped in black foil to avoid distraction by reflection or light effects. Behavioural experiments were conducted for up to five minutes (300 seconds). Before any experimental behaviour assay, the length of *D. pugilator* shells and the width of *C. maenas* carapaces were measured. Due to practical limitations, *H. diversicolor* specimens were not weighed. In addition, the sex of *C. maenas* was noted based on their abdomen shape with females being selected for the experiment, whereas this was not possible for *D. pugilator* and *H. diversicolor*, for which both sexes were used.

The feeding cue was prepared by mashing five adult blue mussels (*Mytilus edulis*) and diluting one volume of mussel homogenate in nine volumes of seawater at pH = 8.2.

Either *D. pugilator* and *C. maenas* specimens were randomly drawn from acclimatised populations raised in pH-controlled communal tanks (pH = 8.2) and held in the middle of the testing test tank using plastic pipe rings. Next, 0.1 mL ten-times diluted mussel juice was randomly injected using a graduated plastic syringe at either side of the tank into an approx. 1 $cm^3$ yellow synthetic sponge ballasted inside a small metal screw nut. The feeding cue was allowed to diffuse in the seawater for 10 s before the crab was released by lifting the pipe ring. Sets of nine *H. diversicolor* were carefully placed from a holding tub into the centre of each of nine test dishes holding conditioned water, and time to burrow the entire head was measured individually using a stopwatch “laps” function.

<u>Details of respiratory rates measurements</u>

Oxygen levels were recorded in *C. maenas* using a fibre-optic oxygen sensor (OXSP5, Pyroscience GmbH, Germany) plugged into FireStingO2 oxygen meter (FSO2-4, Pyroscience GmbH, Germany) and read by Pyro Oxygen Logger software (v 3.315 (c) 2019, Pyroscience GmbH, Germany). Sensors were calibrated for 0% oxygen using factory-calibration and for 100% oxygen using over-oxygenated seawater at pH = 8.2. Respiration rates of *C. maenas* were measured in sealed glass jars containing the same conditioned water in which the behavioural assay was performed beforehand. Oxygen measurements were conducted for 300 seconds. After the first 120 s, mussel juice was added to the seawater. Because no notable changes in oxygen consumption were noted after addition of feeding cue, the entire 300 seconds were kept for statistical analysis, with the exception of clear outliers during the first 10 s that were discarded due to the time the sensor needed to recalibrate after placing an individual into the jar. Once an animal was used for the respirometry assay, it was allowed to recover in normal pH control condition in community tanks. The raw oxygen measurements are available as electronic supplementary material.

<u>Details of statistical analyses</u>

*Factorial design of pH drop, stress metabolites, and donor*

The relationship between oxygen levels and time (for 300 seconds) was modelled using linear models (see R code for details). The slope, Pearson’s r and P were extracted from the model. All models were significant with P ranging from 0.01 to $< 2.2E^{-16}$. Next, the oxygen consumption was transformed using the additive-inverse slope coefficient standardised to the carapace width and rescaled to the mean oxygen consumption of the control CM to yield relative oxygen consumption per second per size, or respiration rate. Respiration rate data was transformed by Yeo-Johnson normalisation from the *BestNormalize* R package (Peterson and Cavanaugh, 2020) ($\lambda = -0.558$, model residuals: Shapiro-Wilk’s W = 0.99, P = 0.45, Bartlett’s K-squared = 0.61, P = 0.89). Effect size thresholds followed the classification given in Sawilowsky (2009): $|d| > 0.01$: very small, $|d| > 0.2$: small, $|d| > 0.5$: medium, $|d| > 0.8$: large, $|d| > 1.20$: very large, $|d| > 2.0$: huge.

*Additional comparisons between CM100% and CM or SM*

In addition to the full factorial design, two separate conditions were tested to better understand the effect of metabolites and discriminate the involvement of pH drop-induced stress metabolites *versus* higher concentrations of control metabolites induced by a higher metabolism in pH drop-stressed specimens. Therefore, the effects of undiluted control metabolites from *S. aurata* (CM100%) on the feeding response of *D. pugilator* and the burrowing behaviour of *H. diversicolor* were compared to the effects of ten-time diluted treatments of stress metabolites SM and control metabolites CM (from the general factorial design). False discovery rate-corrected pairwise comparisons between SM, CM100% and CM were performed using the estimated means (*emmeans*) R package (Lenth, 2019) after either generalised linear models for logit regression with binomial distribution (escaping response in *D. pugilator* and avoidance response in *H. diversicolor*), or Cox proportional hazard models (time-to-success analysis).

**Supplementary Results and Discussion**

Partitioning avoidance responses into escaping and freezing in crabs

Partitioning the avoidance responses of crabs into freezing or escaping patterns allowed a better understanding of their locomotion response to predictors (Figures S1-S2). Apart from *D. pugilator* freezing significantly less in presence of conspecific metabolites (24%) than of *S. aurata* metabolites (53%, Z = 2.46, P = 0.0139), no significant patterns were observed in the freezing responses of *D. pugilator* (Tables S4-S5) and *C. maenas* (Tables S10-S11) to pH drop and stress metabolite predictors. Instead, observed changes in avoidance patterns were mainly attributed to escaping responses. Escaping patterns of *D. pugilator* did not vary with the pH drop, stress metabolites, nor donor predictors (Table S6) which was also true for specimens that received conspecific metabolites but not *S. aurata* metabolites. Indeed, pairwise comparisons revealed that *S. aurata* control metabolites in pH drop induced a significant eightfold increase in escaping (40% in pH drop treatment) in *D. pugilator* compared to control metabolites in control pH (5% in CM treatment, Z = -3.17, P = 0.0091, Table S8). Furthermore, control metabolites from *S. aurata* augmented *D. pugilator* escaping in pH drop (Z = -3.09, P = 0.0079) but not in control pH (P = 0.9876), compared to control metabolites from conspecifics. Stress metabolites from *S. aurata* also significantly increased the avoidance of *D. pugilator* compared to conspecific stress metabolites (Z = -2.35, P = 0.0380). A similar pattern was observed on the escaping response of *C. maenas* wherein effects of pH drop, stress metabolite, and donor predictors were not significant overall (which held true in specimens exposed to conspecific metabolites) but the metabolite:donor interaction term was significant (Z = 2.06, P = 0.0395, Tables S12-S13, Figures S1-S2). However, both pH drop (81%, Z = -3.49, P = 0.0029) and SM (60%, Z = -2.82, P = 0.0096) treatments significantly increased the display of escaping compared to the control CM (6%) in *C. maenas* exposed to *S. aurata* metabolites. The escaping response of *C. maenas* to stress metabolites dropped to 28% in the pH drop+SM treatment compared to the pH drop treatment (Z = 2.92, P = 0.0096). Last, stress metabolites from *S. aurata* significantly increased escaping compared to that from conspecifics (Z = -2.98, P = 0.0116).

Confounding effects of covariates

Interestingly, there were slight confounding effects of covariates on avoidance responses which diminished with time as conditioning water was reused in *D. pugilator* (Z = -1.98, P = 0.0480) and *H. diversicolor* (Z = -5.2980, P < 0.0001). Escaping was more pronounced in larger individuals of *D. pugilator* (Z = 2.49, P = 0.0129, Figure S4, Table S8). However, no confounding effects of number of water uses nor crab sizes were observed on the time-to-success responses.

Relationship between avoidance and time-to-success responses

Cox proportional hazard models were used to estimate whether the observed avoidance behaviour altered the success probability of tested animals (Figure S5). Specimens of *D. pugilator* that displayed an avoidance response were associated with lower feeding success ratios compared to those that did not (P < 0.0001). This was also true for specimens of *D. pugilator* that tended to freeze (P < 0.0001) but not for those displaying an escaping response (P = 0.2060). This may be that *D. pugilator* individuals that tried to escape the testing tank went along the walls and eventually found the feeding cue by chance rather than by active foraging (own observations). Avoidance display was also associated with lower feeding success ratios in *C. maenas* (P < 0.0001), which held true for both the escaping (P = 0.0047) and freezing (P = 0.0003) patterns. On the other hand, there was no relationship between the burrowing success ratio and the display of avoidance behaviours in *H. diversicolor.*

Comparison between CM100% and CM or SM

Next, we investigated whether the effects of stress metabolites were associated with more regularly excreted metabolic wastes using behavioural observations from different concentrations of sea bream metabolites (Figure S6). Neither the success ratios nor the percentage of avoidance in *H. diversicolor* and escaping in *D. pugilator* depended on the concentration of sea bream control metabolites as there were no significant differences between CM and CM100% (Tables S17-S18). On the other hand, SM significantly lowered burrowing success ratios in *H. diversicolor* (P = 0.0266) and increased avoidance in *H. diversicolor* (Z = -5.88, P < 0.0001) and escaping in *D. pugilator* (Z = 2.27, P = 0.023) compared to CM100% treatment.

**Supplementary Figures**

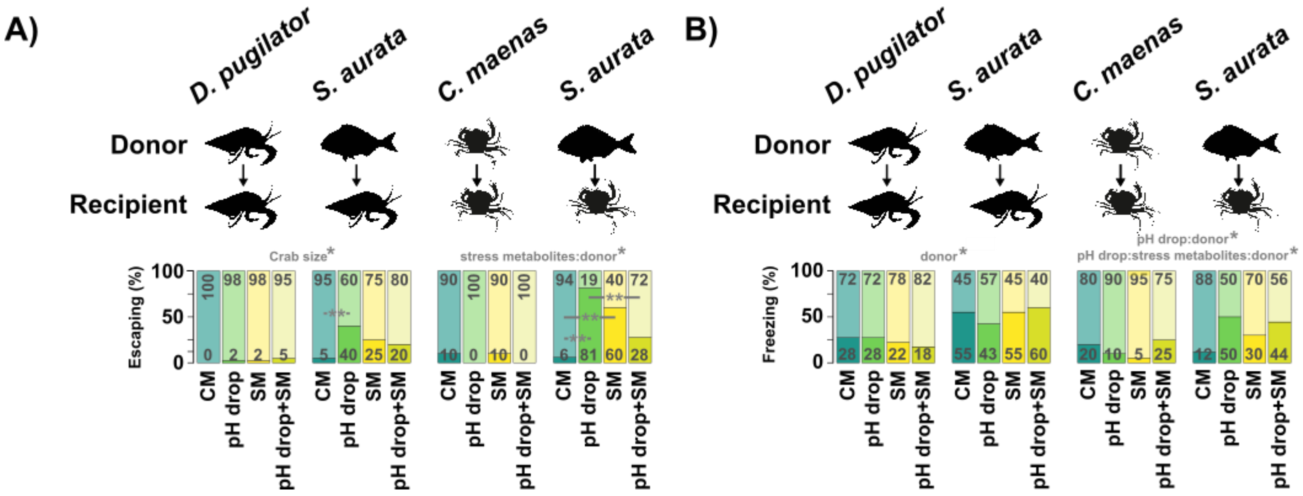


**Figure S1**. Percentage of escaping (A and B) and freezing (C and D) behaviour in small hermit crab *Diogenes pugilator* and green shore crab *Carcinus maenas* depending on main predictors (pH drop, stress metabolites, and donor). Metabolite donors (conspecifics, or the heterospecific gilt-head sea bream *Sparus aurata*) were conditioned in control pH (pH = 8.2) or pH drop (pH = 7.6) to release control metabolites or stress metabolites, respectively. Recipient species were exposed to conspecific or heterospecific metabolites in both control pH and pH drop. Split bars represent the presence (dark area) or the absence (light area) of escaping and freezing behaviours. Significant main predictors, their interaction terms, and covariates (number of water uses, crab size) are shown above plots. Significant pairwise comparisons between treatments are shown below as horizontal black bars. *: $P \leq 0.05$, **: $P < 0.01$, ***: $P < 0.001$, ****: $P < 0.0001$. Significance was retrieved from binomial generalised linear models. CM: control metabolites in control pH, pH drop: control metabolites in pH drop, SM: stress metabolites in control pH, pH drop+SM: stress metabolites in pH drop.

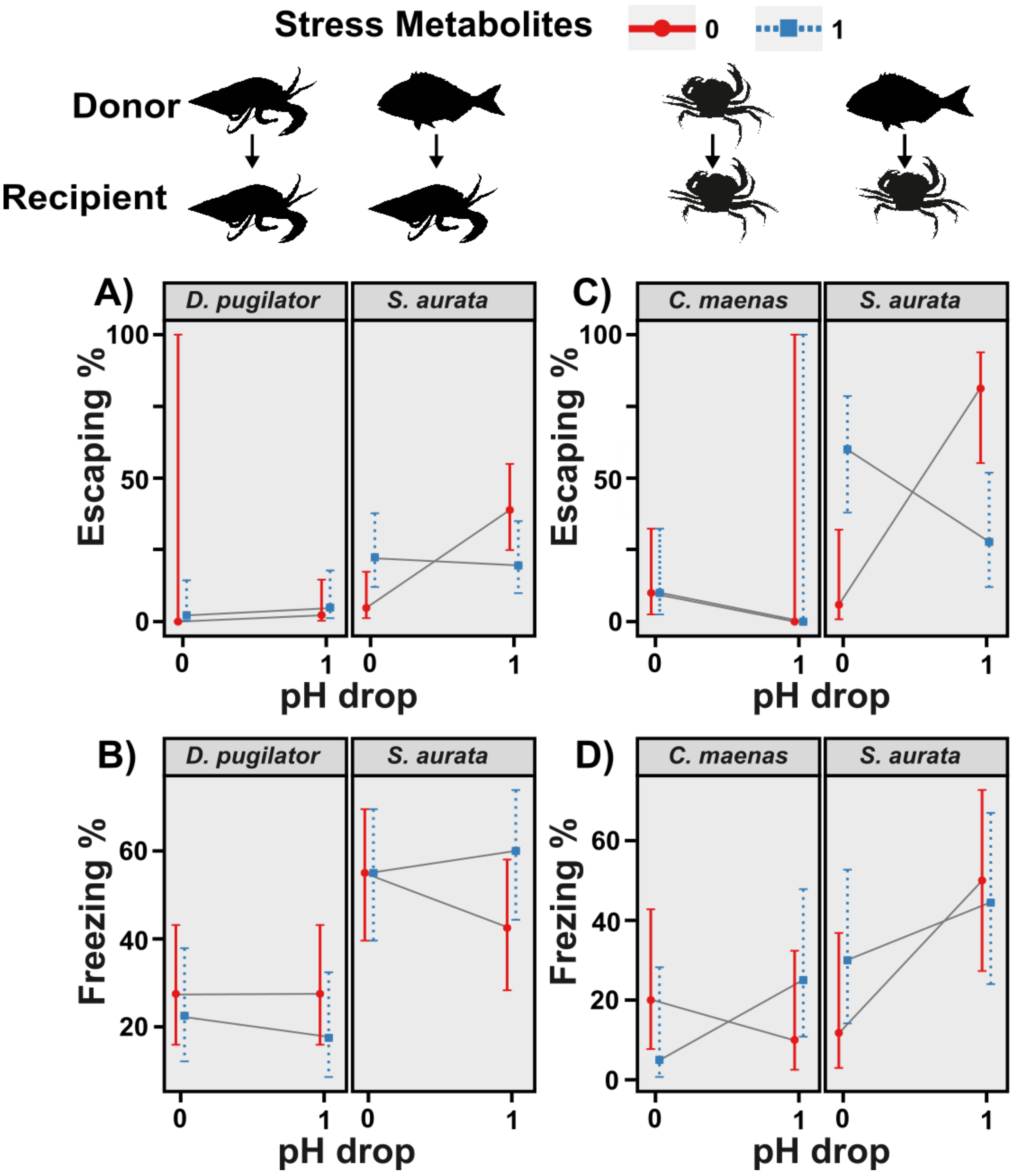


**Figure S2**. Predicted escaping (A and C) and freezing (B and D) percentages showing the marginal effects of three predictors (pH drop, stress metabolites, donor) in small hermit crab *Diogenes pugilator* (left, A and B), green shore crab *Carcinus maenas* (right, C and D). Metabolite donors (conspecifics, or the heterospecific gilt-head sea bream *Sparus aurata*) were conditioned in control pH (pH = 8.2) or pH drop (pH = 7.6) to trigger the release of control metabolites or stress metabolites, respectively. Recipient species were exposed to conspecific or heterospecific metabolites in both control pH and pH drop. Predictors are binary coded as 0 (control metabolites, control pH) and 1 (stress metabolites, pH drop). Crossing solid grey lines represent an interacting effect of pH and metabolites for each donor.

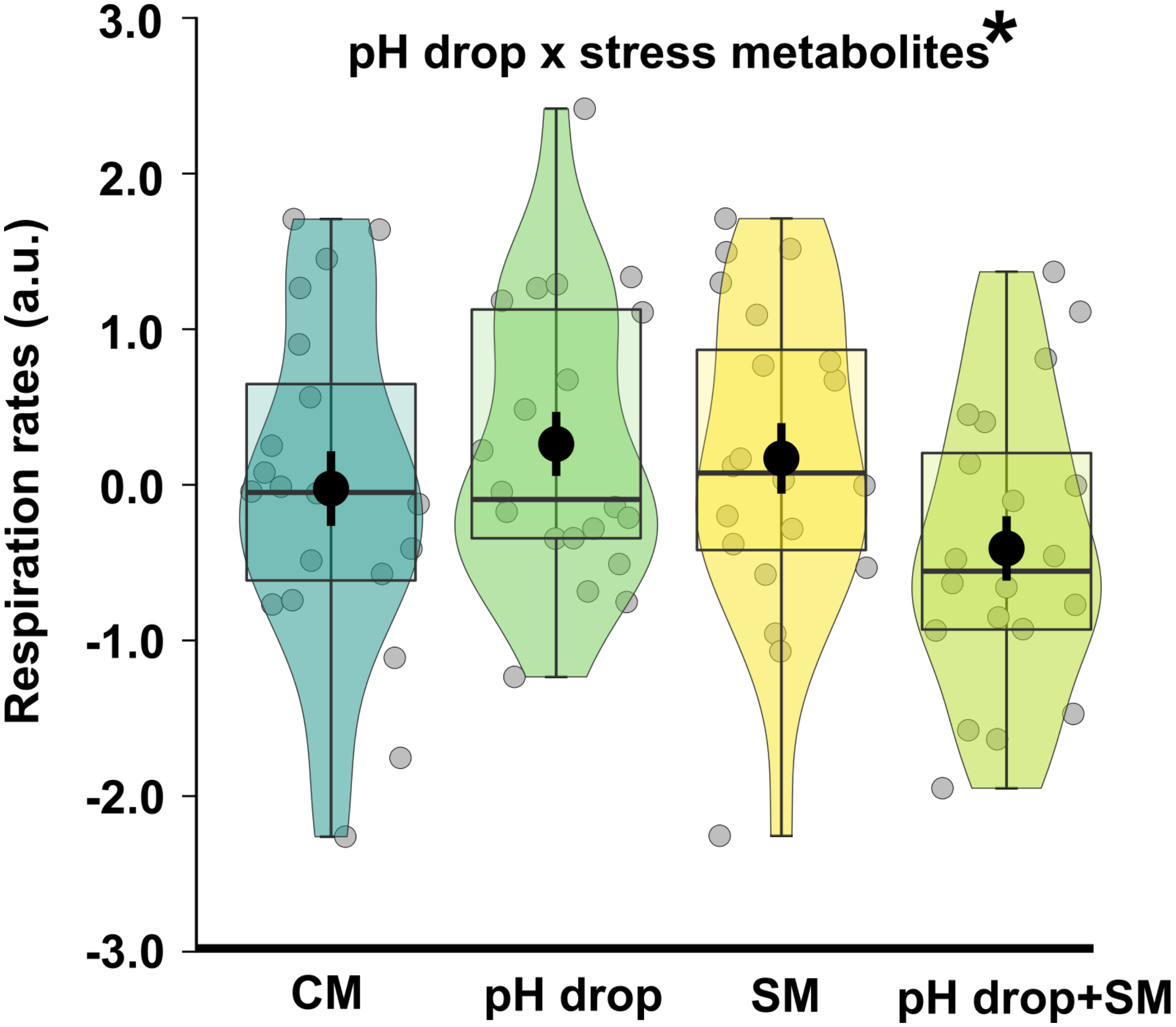


**Figure S3.** Effects of treatments on respiration rate of green shore crab *Carcinus maenas*. Conspecific metabolite donors were conditioned in control pH (pH = 8.2) or pH drop (pH = 7.6) to release control metabolites (CM) or stress metabolites (SM), respectively. *C. maenas* were exposed to metabolites in both control pH and pH drop. Respiration rate is displayed as the standardised, rescaled slope coefficient of oxygen levels over time. Boxes represent median, 25%-75% quartiles, and whiskers are minimum and maximum values within 1.5 IQR (interquartile range). Jittered transformed data given as open circles. Black dots and thick vertical black bars represent the mean ± standard error values of each treatment. Significant factors (pH drop, stress metabolites) are shown above plots. Pairwise comparisons between treatments were not significant. CM: control metabolites in control pH, pH drop: control metabolites in pH drop, SM: stress metabolites in control pH, pH drop+SM: stress metabolites in pH drop.

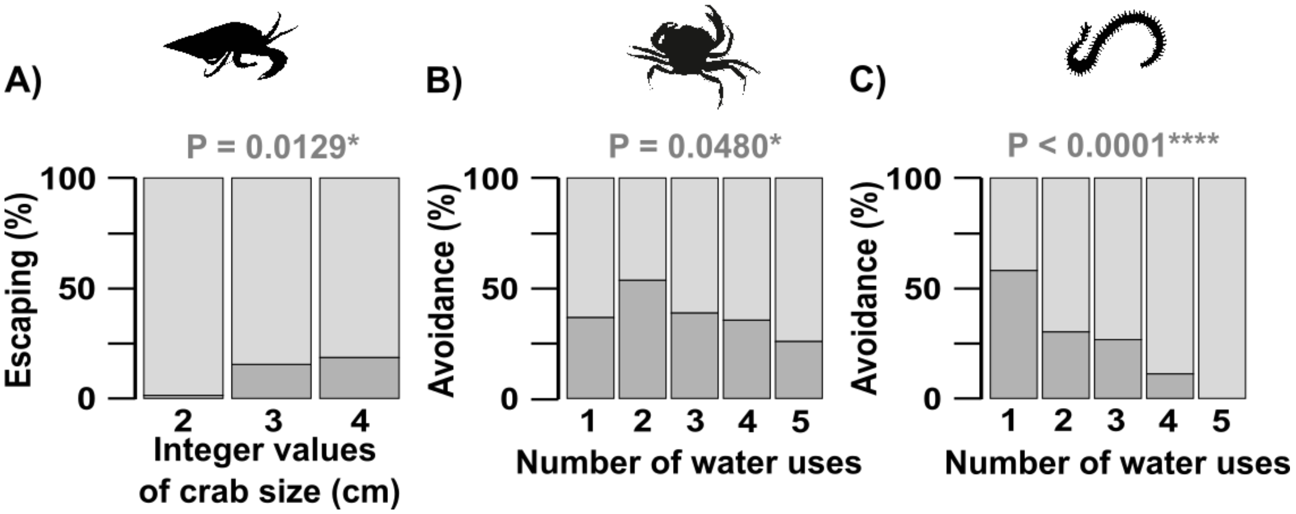


**Figure S4.** Effect of confounding variables significantly altering avoidance behaviour. A) Escaping behaviour of small hermit crab *Diogenes pugilator* significantly reduced with crab size (shown here as integer values to represent a continuous range of 2.0 cm to 4.7 cm. The reuse of conditioning seawater significantly decreased avoidance behaviour (including freezing and escaping) of green shore crab *Carcinus maenas* (B) and avoidance behaviour (including freezing, curling, flipping, and slime secretion) of harbour ragworm *Hediste diversicolor* (C). Split bars represent the presence (dark area) or the absence (light area) of observed behaviours. Significance was retrieved from binomial generalised linear models.

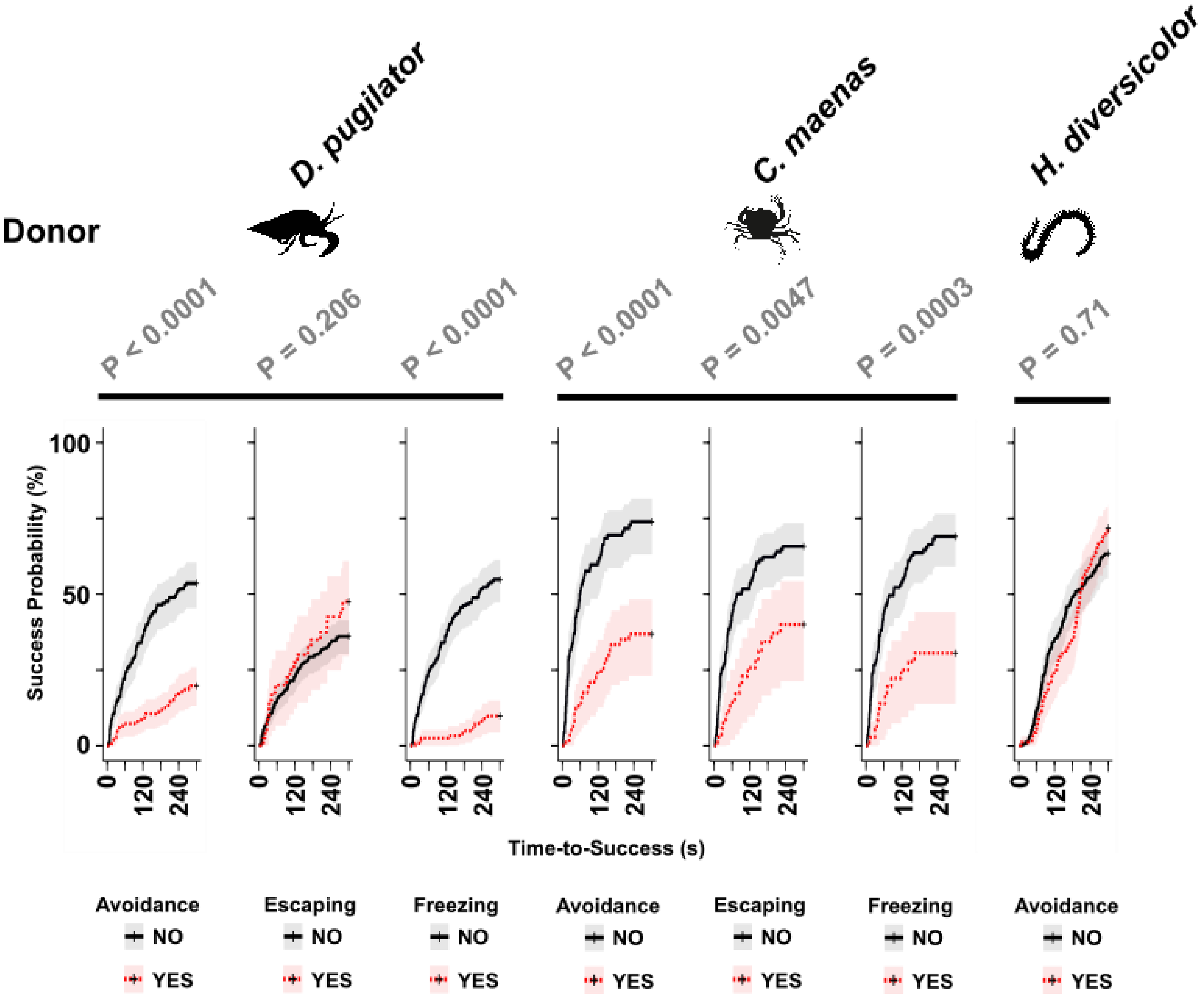


**Figure S5.** Kaplan-Meier curves of time-to-success analysis showing the effects of displayed avoidance behaviour on the success probability (cumulative event) over time in small hermit crab *Diogenes pugilator* (left), green shore crab *Carcinus maenas* (middle), and harbour ragworm *Hediste diversicolor* (right). Avoidance behaviour included freezing and escaping in crabs, and freezing, curling, flipping, and slime secretion in harbour ragworms. Avoidance was also split into freezing and escaping in crabs. P-values represent the significance level comparing the presence vs absence of the studied behaviour on the success ratio according to a Cox proportional hazard model.

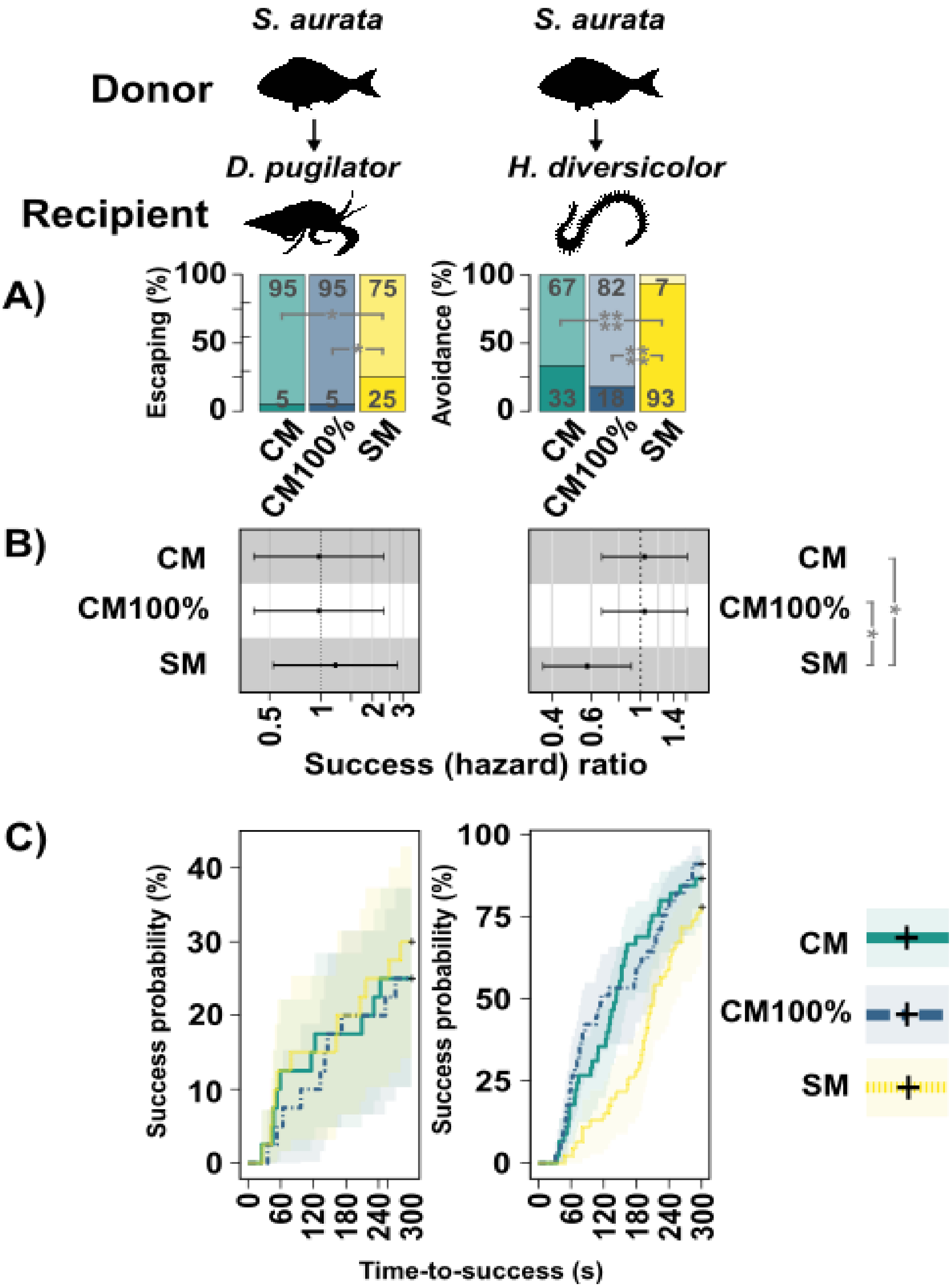


**Figure S6**. Comparison of concentration and type of metabolites originating from gilt-head sea bream *Sparus aurata* on fitness-relevant behaviours of small hermit crab *Diogenes pugilator* (left) and harbour ragworm *Hediste diversicolor* (right). Top panel shows the percent display of escaping behaviour in crabs, and avoidance including freezing, curling, flipping, and slime secretion in harbour ragworms. Split bars represent the presence (dark area) or absence (light area) of escaping or avoidance behaviour. Lower panels show time-to-success analysis (finding the feeding cue in crab or burrowing the entire head in ragworm). Thick lines (with confidence interval as shaded areas) show the probability of success in function of time over 300 seconds. CM: 10-times diluted sea bream control metabolites, CM100%: undiluted sea bream control metabolites. SM: 10-times diluted sea bream stress metabolites. Pairwise comparisons between CM100% and either CM or SM are represented with grey lines. *: $P \leq 0.05$, **: $P < 0.01$, ***: $P < 0.001$, ****: $P < 0.0001$. P-values represent the significance level comparing the presence vs absence of the studied behaviour on the success ratio according to Cox proportional hazard models.

### Supplementary Tables

**Table S1.** Summary of conditions for behavioural assays. Tested species were: small hermit crab *Diogenes pugilator*, green shore crab *Carcinus maenas*, harbour ragworm *Hediste diversicolor*, and gilt-head sea bream *Sparus aurata*. Factors were: pH drop (0 for control pH, pH = 8.2 *vs.* 1 for pH drop, pH = 7.6), stress metabolites (0 for control metabolites putatively released at pH = 8.2 *vs.* 1 for stress metabolites putatively released at pH = 7.6), and donor (conspecific, or heterospecific i.e. *S. aurata*. Conditioned water from *S. aurata* (except stated otherwise) and *H. diversicolor* was tenfold diluted in the system water at the desired pH. *n* represents the sample size.

| Treatment | Donor | Recipient | Donor factor | pH | Metabolites | pH adjustment | Dilution | $pCO_2$ | *n* |
|---|---|---|---|---|---|---|---|---|---|
| **Factorial design** | | | | | | | | | |
| CM | *D. pugilator* | *D. pugilator* | Conspecific | 0 | 0 | - | - | 400 | 40 |
| pH drop | *D. pugilator* | *D. pugilator* | Conspecific | 1 | 0 | HCl addition to CM conditioned water | - | 400 | 40 |
| SM | *D. pugilator* | *D. pugilator* | Conspecific | 0 | 1 | Reef buffer addition to SM conditioned water | - | 700 | 40 |
| pH drop+SM | *D. pugilator* | *D. pugilator* | Conspecific | 1 | 1 | - | - | 700 | 40 |
| CM | *S. aurata* | *D. pugilator* | Heterospecific | 0 | 0 | CM conditioned water diluted in pH = 8.2 system water | 10% | 400 | 40 |
| pH drop | *S. aurata* | *D. pugilator* | Heterospecific | 1 | 0 | CM conditioned water diluted in pH = 7.6 system water | 10% | 670 | 40 |
| SM | *S. aurata* | *D. pugilator* | Heterospecific | 0 | 1 | SM conditioned water diluted in pH = 8.2 system water | 10% | 430 | 40 |
| pH drop+SM | *S. aurata* | *D. pugilator* | Heterospecific | 1 | 1 | SM conditioned water diluted in pH = 7.6 system water | 10% | 700 | 40 |
| CM | *C. maenas* | *C. maenas* | Conspecific | 0 | 0 | - | - | 400 | 29 |
| pH drop | *C. maenas* | *C. maenas* | Conspecific | 1 | 0 | HCl addition to CM conditioned water | - | 400 | 20 |
| SM | *C. maenas* | *C. maenas* | Conspecific | 0 | 1 | Reef buffer addition to SM conditioned water | - | 700 | 35 |
| pH drop+SM | *C. maenas* | *C. maenas* | Conspecific | 1 | 1 | - | - | 700 | 35 |
| CM | *S. aurata* | *C. maenas* | Heterospecific | 0 | 0 | CM conditioned water diluted in pH = 8.2 system water | 10% | 400 | 16 |
| pH drop | *S. aurata* | *C. maenas* | Heterospecific | 1 | 0 | CM conditioned water diluted in pH = 7.6 system water | 10% | 670 | 17 |
| SM | *S. aurata* | *C. maenas* | Heterospecific | 0 | 1 | SM conditioned water diluted in pH = 8.2 system water | 10% | 430 | 20 |
| pH drop+SM | *S. aurata* | *C. maenas* | Heterospecific | 1 | 1 | SM conditioned water diluted in pH = 7.6 system water | 10% | 700 | 18 |
| CM | *H. diversicolor* | *H. diversicolor* | Conspecific | 0 | 0 | CM conditioned water diluted in pH = 8.2 system water | 10% | 400 | 45 |
| pH drop | *H. diversicolor* | *H. diversicolor* | Conspecific | 1 | 0 | CM conditioned water diluted in pH = 7.6 system water | 10% | 670 | 27 |
| SM | *H. diversicolor* | *H. diversicolor* | Conspecific | 0 | 1 | SM conditioned water diluted in pH = 8.2 system water | 10% | 430 | 27 |
| pH drop+SM | *H. diversicolor* | *H. diversicolor* | Conspecific | 1 | 1 | SM conditioned water diluted in pH = 7.6 system water | 10% | 700 | 45 |
| CM | *S. aurata* | *H. diversicolor* | Heterospecific | 0 | 0 | CM conditioned water diluted in pH = 8.2 system water | 10% | 400 | 45 |
| pH drop | *S. aurata* | *H. diversicolor* | Heterospecific | 1 | 0 | CM conditioned water diluted in pH = 7.6 system water | 10% | 670 | 45 |
| SM | *S. aurata* | *H. diversicolor* | Heterospecific | 0 | 1 | SM conditioned water diluted in pH = 8.2 system water | 10% | 430 | 46 |
| pH drop+SM | *S. aurata* | *H. diversicolor* | Heterospecific | 1 | 1 | SM conditioned water diluted in pH = 7.6 system water | 10% | 700 | 45 |
| **Additional treatments** | | | | | | | | | |
| CM100% | *S. aurata* | *H. diversicolor* | Heterospecific | 0 | 0 | - | - | 400 | 45 |
| CM100% | *S. aurata* | *D. pugilator* | Heterospecific | 0 | 0 | - | - | 400 | 40 |

**Table S2.** Results of post-hoc pairwise comparisons of interest from the interaction term between the three main predictors (pH drop, stress metabolites, donor) of the Cox proportional hazard model on the time-to-success analysis in small hermit crab (*Diogenes pugilator*) receiving metabolites from conspecifics (*D. pugilator*) or heterospecific gilt-head sea bream (*Sparus aurata*). Post-hoc tests are split into two terms expressed as '(pH drop:stress metabolites)|donor' and 'donor|(pH drop:stress metabolites)', respectively comparing the four treatments (CM, pH drop, SM, and pH drop+SM) within each donor group, and comparing the donor effect within each treatment (CM, pH drop, SM, or pH drop+SM). Events are the success to find a feeding cue in 300 seconds. Unsuccessful animals are censored. Significance ($P \leq 0.05$) is shown by p-values in bold. Results are expressed on the log (not the response) scale. SE: standard error of estimate. P-values are adjusted using false discovery rate adjustment for the number of comparisons within each post-hoc term. Comparisons were calculated using the emmeans R package (Lenth, 2019) on the Cox proportional hazard model computed via the survival R package (Therneau and Grambsch, 2000; Therneau, 2020). CM: control metabolites tested in control pH (pH = 8.2), pH drop: control metabolites tested in pH drop (pH = 7.6), SM: stress metabolites tested in control pH (pH = 8.2), pH drop+SM: stress metabolites tested in pH drop (pH = 7.6).

| Interaction term | Post-hoc Term | Group | Comparison | Estimate | SE | Z | Padj |
|---|---|---|---|---|---|---|---|
| pH drop:stress metabolites:donor | (pH drop:stress metabolites)\|donor | *D. pugilator* | CM - pH drop | 0.7546 | 0.3530 | 2.1378 | 0.1952 |
| pH drop:stress metabolites:donor | (pH drop:stress metabolites)\|donor | *D. pugilator* | CM - SM | 0.2100 | 0.3167 | 0.6631 | 0.6030 |
| pH drop:stress metabolites:donor | (pH drop:stress metabolites)\|donor | *D. pugilator* | CM - pH drop+SM | 0.3836 | 0.3263 | 1.1756 | 0.4710 |
| pH drop:stress metabolites:donor | (pH drop:stress metabolites)\|donor | *D. pugilator* | pH drop - SM | -0.5447 | 0.3601 | -1.5127 | 0.3911 |
| pH drop:stress metabolites:donor | (pH drop:stress metabolites)\|donor | *D. pugilator* | pH drop - pH drop+SM | -0.3710 | 0.3685 | -1.0068 | 0.4710 |
| pH drop:stress metabolites:donor | (pH drop:stress metabolites)\|donor | *D. pugilator* | SM - pH drop+SM | 0.1736 | 0.3339 | 0.5201 | 0.6030 |
| pH drop:stress metabolites:donor | (pH drop:stress metabolites)\|donor | *S. aurata* | CM - pH drop | -0.4345 | 0.4141 | -1.0493 | 0.7898 |
| pH drop:stress metabolites:donor | (pH drop:stress metabolites)\|donor | *S. aurata* | CM - SM | -0.1934 | 0.4282 | -0.4518 | 0.7898 |
| pH drop:stress metabolites:donor | (pH drop:stress metabolites)\|donor | *S. aurata* | CM - pH drop+SM | -0.3338 | 0.4141 | -0.8060 | 0.7898 |
| pH drop:stress metabolites:donor | (pH drop:stress metabolites)\|donor | *S. aurata* | pH drop - SM | 0.2411 | 0.3934 | 0.6127 | 0.7898 |
| pH drop:stress metabolites:donor | (pH drop:stress metabolites)\|donor | *S. aurata* | pH drop - pH drop+SM | 0.1007 | 0.3780 | 0.2665 | 0.7898 |
| pH drop:stress metabolites:donor | (pH drop:stress metabolites)\|donor | *S. aurata* | SM - pH drop+SM | -0.1403 | 0.3934 | -0.3566 | 0.7898 |
| pH drop:stress metabolites:donor | donor\|(pH drop: stress metabolites) | CM | *S. aurata - D. pugilator* | 1.0613 | 0.3843 | **2.7613** | **0.0230** |
| pH drop:stress metabolites:donor | donor\|(pH drop: stress metabolites) | pH drop | *S. aurata - D. pugilator* | -0.1279 | 0.3852 | -0.3320 | 0.7399 |
| pH drop:stress metabolites:donor | donor\|(pH drop: stress metabolites) | SM | *S. aurata - D. pugilator* | 0.6578 | 0.3689 | 1.7833 | 0.1491 |
| pH drop:stress metabolites:donor | donor\|(pH drop: stress metabolites) | pH drop+SM | *S. aurata - D. pugilator* | 0.3439 | 0.3610 | 0.9527 | 0.4544 |

**Table S3**. Results of post-hoc pairwise comparisons of interest from the interaction term between the three main predictors (pH drop, stress metabolites, donor) of the binomial generalised linear model on the avoidance behaviour (including freezing and escaping) in small hermit crab (*Diogenes pugilator*) receiving metabolites from conspecifics (*D. pugilator*) or heterospecific gilt-head sea bream (*Sparus aurata*). Post-hoc tests are split into two terms expressed as ‘(pH drop:stress metabolites)|donor’ and ‘donor|(pH drop:stress metabolites)’, respectively comparing the four treatments (CM, pH drop, SM, and pH drop+SM) within each donor group, and comparing the donor effect within each treatment (CM, pH drop, SM, or pH drop+SM). Significance ($P \leq 0.05$) is shown by estimates and p-values in bold. Results are expressed on the log (not the response) scale. SE: standard error of estimate. P-values are adjusted using false discovery rate adjustment for the number of comparisons within each post-hoc term. Comparisons were calculated using the emmeans R package (Lenth, 2019) on the generalised linear model computed via the stats R package (R Core Team, 2020). CM: control metabolites tested in control pH (pH = 8.2), pH drop: control metabolites tested in pH drop (pH = 7.6), SM: stress metabolites tested in control pH (pH = 8.2), pH drop+SM: stress metabolites tested in pH drop (pH = 7.6).

| Model term | Post-hoc Term | Group | Comparison | Estimate | SE | Z | Padj |
|---|---|---|---|---|---|---|---|
| pH drop:stress metabolites:donor | (pH drop:stress metabolites)\|donor | *D. pugilator* | CM - pH drop | -0.1221 | 0.4944 | -0.2470 | 0.8049 |
| pH drop:stress metabolites:donor | (pH drop:stress metabolites)\|donor | *D. pugilator* | CM - SM | 0.1292 | 0.5086 | 0.2540 | 0.8049 |
| pH drop:stress metabolites:donor | (pH drop:stress metabolites)\|donor | *D. pugilator* | CM - pH drop+SM | 0.2674 | 0.5184 | 0.5157 | 0.8049 |
| pH drop:stress metabolites:donor | (pH drop:stress metabolites)\|donor | *D. pugilator* | pH drop - SM | 0.2513 | 0.5024 | 0.5003 | 0.8049 |
| pH drop:stress metabolites:donor | (pH drop:stress metabolites)\|donor | *D. pugilator* | pH drop - pH drop+SM | 0.3895 | 0.5123 | 0.7603 | 0.8049 |
| pH drop:stress metabolites:donor | (pH drop:stress metabolites)\|donor | *D. pugilator* | SM - pH drop+SM | 0.1382 | 0.5260 | 0.2626 | 0.8049 |
| pH drop:stress metabolites:donor | (pH drop:stress metabolites)\|donor | *S. aurata* | CM - pH drop | -0.8979 | 0.4841 | -1.8549 | 0.1908 |
| pH drop:stress metabolites:donor | (pH drop:stress metabolites)\|donor | *S. aurata* | CM - SM | -0.89879 | 0.4841 | -1.8549 | 0.1908 |
| pH drop:stress metabolites:donor | (pH drop:stress metabolites)\|donor | *S. aurata* | CM - pH drop+SM | -0.6466 | 0.4691 | -1.3784 | 0.3361 |
| pH drop:stress metabolites:donor | (pH drop:stress metabolites)\|donor | *S. aurata* | pH drop - SM | 0.0000 | 0.5164 | 0.0000 | 1.0000 |
| pH drop:stress metabolites:donor | (pH drop:stress metabolites)\|donor | *S. aurata* | pH drop - pH drop+SM | 0.2513 | 0.5024 | 0.5003 | 0.7403 |
| pH drop:stress metabolites:donor | (pH drop:stress metabolites)\|donor | *S. aurata* | SM - pH drop+SM | 0.2513 | 0.5024 | 0.5003 | 0.7403 |
| pH drop:stress metabolites:donor | donor\|(pH drop: stress metabolites) | CM | *S. aurata - D. pugilator* | -1.1701 | 0.4758 | **-2.4591** | **0.0139** |
| pH drop:stress metabolites:donor | donor\|(pH drop: stress metabolites) | pH drop | *S. aurata - D. pugilator* | -1.9459 | 0.5024 | **-3.8734** | **0.0001** |
| pH drop:stress metabolites:donor | donor\|(pH drop: stress metabolites) | SM | *S. aurata - D. pugilator* | -2.1972 | 0.5164 | **-4.2549** | **0.0001** |
| pH drop:stress metabolites:donor | donor\|(pH drop: stress metabolites) | pH drop+SM | *S. aurata - D. pugilator* | -2.0841 | 0.5123 | **-4.0683** | **0.0001** |

**Table S4**. Results of binomial generalised linear model for the main effects of predictors (pH drop, stress metabolites, donor) on the freezing behaviour in small hermit crab (*Diogenes pugilator*). Significance ($P \leq 0.05$) of predictors is shown in italics with the p-values and estimates in bold. Sample size: 320 observations. Overall significance of models from Chi-squared analyses of deviance when including only predictors was $P < 0.0001$. Covariates were dropped from the model after an analysis of deviance showed that they passed the Chi-squared test (number of water uses: $P = 0.4922$, crab size: $P = 0.4224$). SE is the standard error of estimate. Modelled using the stats R package (R Core Team, 2020).

| **Predictor** | **Estimate** | **SE** | **Z** | **P** |
|---|---|---|---|---|
| (Intercept) | -0.9694 | 0.3541 | -2.7376 | 0.0062 |
| pH drop | 0.0000 | 0.5008 | 0.0000 | 1.0000 |
| stress metabolites | -0.2674 | 0.5184 | -0.5157 | 0.6060 |
| donor | 1.1701 | 0.4758 | **2.4591** | **0.0139** |
| pH drop:stress metabolites | -0.3138 | 0.7532 | -0.4167 | 0.6769 |
| pH:donor | -0.5030 | 0.6739 | -0.7464 | 0.4554 |
| stress metabolites:donor | 0.2674 | 0.6861 | 0.3897 | 0.6968 |
| pH:stress metabolites:donor | 1.0216 | 0.9878 | 1.0342 | 0.3011 |

**Table S5**. Results of post-hoc pairwise comparisons of interest from the interaction term between the three main predictors (pH drop, stress metabolites, donor) of the binomial generalised linear model on the freezing behaviour in small hermit crab (*Diogenes pugilator*) receiving metabolites from conspecifics (*D. pugilator*) or heterospecific gilt-head sea bream (*Sparus aurata*). Post-hoc tests are split into two terms expressed as '(pH drop:stress metabolites)|donor' and 'donor|(pH drop:stress metabolites)', respectively comparing the four treatments (CM, pH drop, SM, and pH drop+SM) within each donor group, and comparing the donor effect within each treatment (CM, pH drop, SM, or pH drop+SM). Significance ($P \leq 0.05$) is shown by p-values in bold. Results are expressed on the log (not the response) scale. SE: standard error of estimate. P-values are adjusted using false discovery rate adjustment for the number of comparisons within each post-hoc term. Comparisons were calculated using the emmeans R package (Lenth, 2019) on the generalised linear model computed via the stats R package (R Core Team, 2020). CM: control metabolites tested in control pH (pH = 8.2), pH drop: control metabolites tested in pH drop (pH = 7.6), SM: stress metabolites tested in control pH (pH = 8.2), pH drop+SM: stress metabolites tested in pH drop (pH = 7.6).

| **Model term** | **Post-hoc Term** | **Group** | **Comparison** | **Estimate** | **SE** | **Z** | **Padj** |
|---|---|---|---|---|---|---|---|
| pH drop:stress metabolites:donor | (pH drop:stress metabolites)\|donor | *D. pugilator* | CM - pH drop | 0.0000 | 0.5008 | 0.0000 | 1.0000 |
| pH drop:stress metabolites:donor | (pH drop:stress metabolites)\|donor | *D. pugilator* | CM - SM | 0.2674 | 0.5184 | 0.5157 | 0.7273 |
| pH drop:stress metabolites:donor | (pH drop:stress metabolites)\|donor | *D. pugilator* | CM - pH drop+SM | 0.5812 | 0.5464 | 1.0637 | 0.7273 |
| pH drop:stress metabolites:donor | (pH drop:stress metabolites)\|donor | *D. pugilator* | pH drop - SM | 0.2674 | 0.5184 | 0.5157 | 0.7273 |
| pH drop:stress metabolites:donor | (pH drop:stress metabolites)\|donor | *D. pugilator* | pH drop - pH drop+SM | 0.5812 | 0.5464 | 1.0637 | 0.7273 |
| pH drop:stress metabolites:donor | (pH drop:stress metabolites)\|donor | *D. pugilator* | SM - pH drop+SM | 0.3138 | 0.5626 | 0.5578 | 0.7273 |
| pH drop:stress metabolites:donor | (pH drop:stress metabolites)\|donor | *S. aurata* | CM - pH drop | 0.5030 | 0.4509 | 1.1154 | 0.5293 |
| pH drop:stress metabolites:donor | (pH drop:stress metabolites)\|donor | *S. aurata* | CM - SM | 0.0000 | 0.4495 | 0.0000 | 1.0000 |
| pH drop:stress metabolites:donor | (pH drop:stress metabolites)\|donor | *S. aurata* | CM - pH drop+SM | -0.2048 | 0.4530 | -0.4521 | 0.7814 |
| pH drop:stress metabolites:donor | (pH drop:stress metabolites)\|donor | *S. aurata* | pH drop - SM | -0.5030 | 0.4509 | -1.1154 | 0.5293 |
| pH drop:stress metabolites:donor | (pH drop:stress metabolites)\|donor | *S. aurata* | pH drop - pH drop+SM | -0.7077 | 0.4544 | -1.5576 | 0.5293 |
| pH drop:stress metabolites:donor | (pH drop:stress metabolites)\|donor | *S. aurata* | SM - pH drop+SM | -0.2048 | 0.4530 | -0.4521 | 0.7814 |
| pH drop:stress metabolites:donor | donor\|(pH drop: stress metabolites) | CM | *S. aurata - D. pugilator* | -1.1701 | 0.4758 | **-2.4591** | **0.0186** |
| pH drop:stress metabolites:donor | donor\|(pH drop: stress metabolites) | pH drop | *S. aurata - D. pugilator* | -0.6671 | 0.4772 | -1.3981 | 0.1621 |
| pH drop:stress metabolites:donor | donor\|(pH drop: stress metabolites) | SM | *S. aurata - D. pugilator* | -1.4374 | 0.4943 | **-2.9077** | **0.0073** |
| pH drop:stress metabolites:donor | donor\|(pH drop: stress metabolites) | pH drop+SM | *S. aurata - D. pugilator* | -1.9561 | 0.5266 | **-3.7144** | **0.0008** |

**Table S6**. Results of binomial generalised linear model for the main effects of predictors (pH drop, stress metabolites, donor) on the escaping behaviour in small hermit crab (*Diogenes pugilator*). Significance ($P \leq 0.05$) is shown by p-values in bold. Sample size: 320 observations. Overall significance of models from Chi-squared analyses of deviance when including only predictors was $P < 0.0001$. Covariates were dropped from the model after an analysis of deviance showed that they passed the Chi-squared test (number of water uses: $P = 0.06588$). SE is the standard error of estimate. Modelled using the stats R package (R Core Team, 2020).

| Predictor | Estimate | SE | Z | P |
|---|---|---|---|---|
| (Intercept) | -22.2371 | 1017.9837 | -0.0218 | 0.9826 |
| pH drop | 14.9973 | 1017.9831 | 0.0147 | 0.9882 |
| stress metabolites | 14.9756 | 1017.9831 | 0.0147 | 0.9883 |
| donor | 15.7622 | 1017.9828 | 0.0155 | 0.9876 |
| crab size | 1.0409 | 0.4188 | **2.4854** | **0.0129** |
| pH drop:stress metabolites | -14.1749 | 1017.9839 | -0.0139 | 0.9889 |
| pH drop:donor | -12.4512 | 1017.9834 | -0.0122 | 0.9902 |
| stress metabolites:donor | -13.2210 | 1017.9834 | -0.0130 | 0.9896 |
| pH drop:stress metabolites:donor | 11.4587 | 1017.9843 | 0.0113 | 0.9910 |

**Table S7**. Results of post-hoc pairwise comparisons of interest from the interaction term between the three main predictors (pH drop, stress metabolites, donor) of the binomial generalised linear model on the escaping behaviour in small hermit crab (*Diogenes pugilator*) receiving metabolites from conspecifics (*D. pugilator*) or heterospecific gilt-head sea bream (*Sparus aurata*). Post-hoc tests are split into two terms expressed as '(pH drop:stress metabolites)|donor' and 'donor|(pH drop:stress metabolites)', respectively comparing the four treatments (CM, pH drop, SM, and pH drop+SM) within each donor group, and comparing the donor effect within each treatment (CM, pH drop, SM, or pH drop+SM). Significance ($P \leq 0.05$) is shown by p-values in bold. Results are expressed on the log (not the response) scale. SE: standard error of estimate. P-values are adjusted using false discovery rate adjustment for the number of comparisons within each post-hoc term. Comparisons were calculated using the emmeans R package (Lenth, 2019) on the generalised linear model computed via the stats R package (R Core Team, 2020). CM: control metabolites tested in control pH (pH = 8.2), pH drop: control metabolites tested in pH drop (pH = 7.6), SM: stress metabolites tested in control pH (pH = 8.2), pH drop+SM: stress metabolites tested in pH drop (pH = 7.6).

| Model term | Post-hoc Term | Subset | Comparison | Estimate | SE | Z | Padj |
|---|---|---|---|---|---|---|---|
| pH drop:stress metabolites:donor | (pH drop:stress metabolites)\|donor | *D. pugilator* | CM - pH drop | -14.9973 | 1017.9831 | -0.0147 | 0.9883 |
| pH drop:stress metabolites:donor | (pH drop:stress metabolites)\|donor | *D. pugilator* | CM - SM | -14.9756 | 1017.9831 | -0.0147 | 0.9883 |
| pH drop:stress metabolites:donor | (pH drop:stress metabolites)\|donor | *D. pugilator* | CM - pH drop+SM | -15.7981 | 1017.9828 | -0.0155 | 0.9883 |
| pH drop:stress metabolites:donor | (pH drop:stress metabolites)\|donor | *D. pugilator* | pH drop - SM | 0.0218 | 1.4376 | 0.0154 | 0.9883 |
| pH drop:stress metabolites:donor | (pH drop:stress metabolites)\|donor | *D. pugilator* | pH drop - pH drop+SM | -0.8007 | 1.2518 | -0.6397 | 0.9883 |
| pH drop:stress metabolites:donor | (pH drop:stress metabolites)\|donor | *D. pugilator* | SM - pH drop+SM | -0.8225 | 1.2535 | -0.6561 | 0.9883 |
| pH drop:stress metabolites:donor | (pH drop:stress metabolites)\|donor | *S. aurata* | CM - pH drop | -2.5461 | 0.8025 | **-3.1726** | **0.0091** |
| pH drop:stress metabolites:donor | (pH drop:stress metabolites)\|donor | *S. aurata* | CM - SM | -1.7546 | 0.8196 | -2.1408 | 0.0969 |
| pH drop:stress metabolites:donor | (pH drop:stress metabolites)\|donor | *S. aurata* | CM - pH drop+SM | -1.5845 | 0.8357 | -1.8982 | 0.0982 |
| pH drop:stress metabolites:donor | (pH drop:stress metabolites)\|donor | *S. aurata* | pH drop - SM | 0.7915 | 0.5003 | 1.5820 | 0.1364 |
| pH drop:stress metabolites:donor | (pH drop:stress metabolites)\|donor | *S. aurata* | pH drop - pH drop+SM | 0.9616 | 0.5220 | 1.8423 | 0.0982 |
| pH drop:stress metabolites:donor | (pH drop:stress metabolites)\|donor | *S. aurata* | SM - pH drop+SM | 0.1701 | 0.5503 | 0.3091 | 0.7573 |
| pH drop:stress metabolites:donor | donor\|(pH drop: stress metabolites) | CM | *S. aurata - D. pugilator* | -15.7622 | 1017.9828 | -0.0155 | 0.9876 |
| pH drop:stress metabolites:donor | donor\|(pH drop: stress metabolites) | pH drop | *S. aurata - D. pugilator* | -3.3110 | 1.0701 | **-3.0940** | **0.0079** |
| pH drop:stress metabolites:donor | donor\|(pH drop: stress metabolites) | SM | *S. aurata - D. pugilator* | -2.5412 | 1.0836 | **-2.3452** | **0.0380** |
| pH drop:stress metabolites:donor | donor\|(pH drop: stress metabolites) | pH drop+SM | *S. aurata - D. pugilator* | -1.5486 | 0.8345 | -1.8545 | 0.0849 |

**Table S8**. Results of post-hoc pairwise comparisons of interest from the interaction term between the three main predictors (pH drop, stress metabolites, donor) of the Cox proportional hazard model on the time-to-success analysis in green shore crab (*Carcinus maenas*) receiving metabolites from conspecifics (*C. maenas*) or heterospecific gilt-head sea bream (*Sparus aurata*). Post-hoc tests are split into two terms expressed as '(pH drop:stress metabolites)|donor' and 'donor|(pH drop:stress metabolites)', respectively comparing the four treatments (CM, pH drop, SM, and pH drop+SM) within each donor group, and comparing the donor effect within each treatment (CM, pH drop, SM, or pH drop+SM). Events are the success to find a feeding cue in 300 seconds. Unsuccessful animals are censored. Significance ($P \leq 0.05$) is shown by p-values in bold. Results are expressed on the log (not the response) scale. SE: standard error of estimate. P-values are adjusted using false discovery rate adjustment for the number of comparisons within each post-hoc term. Comparisons were calculated using the emmeans R package (Lenth, 2019) on the Cox proportional hazard model computed via the survival R package (Therneau and Grambsch, 2000; Therneau, 2020). CM: control metabolites tested in control pH (pH = 8.2), pH drop: control metabolites tested in pH drop (pH = 7.6), SM: stress metabolites tested in control pH (pH = 8.2), pH drop+SM: stress metabolites tested in pH drop (pH = 7.6).

| Interaction term | Post-hoc Term | Group | Comparison | Estimate | SE | Z | Padj |
|---|---|---|---|---|---|---|---|
| pH drop:stress metabolites:donor | (pH drop:stress metabolites)\|donor | *C. maenas* | CM - pH drop | 0.2890 | 0.3620 | 0.7982 | 0.5987 |
| pH drop:stress metabolites:donor | (pH drop:stress metabolites)\|donor | *C. maenas* | CM - SM | -0.1953 | 0.2887 | -0.6762 | 0.5987 |
| pH drop:stress metabolites:donor | (pH drop:stress metabolites)\|donor | *C. maenas* | CM - pH drop+SM | 0.2359 | 0.3089 | 0.7639 | 0.5987 |
| pH drop:stress metabolites:donor | (pH drop:stress metabolites)\|donor | *C. maenas* | pH drop - SM | -0.4842 | 0.3452 | -1.4027 | 0.4821 |
| pH drop:stress metabolites:donor | (pH drop:stress metabolites)\|donor | *C. maenas* | pH drop - pH drop+SM | -0.0530 | 0.3619 | -0.1466 | 0.8835 |
| pH drop:stress metabolites:donor | (pH drop:stress metabolites)\|donor | *C. maenas* | SM - pH drop+SM | 0.4312 | 0.2890 | 1.4922 | 0.4821 |
| pH drop:stress metabolites:donor | (pH drop:stress metabolites)\|donor | *S. aurata* | *CM - pH drop* | 1.5374 | 0.5177 | **2.9694** | **0.0179** |
| pH drop:stress metabolites:donor | (pH drop:stress metabolites)\|donor | *S. aurata* | *CM - SM* | 1.0150 | 0.4092 | **2.4804** | **0.0394** |
| pH drop:stress metabolites:donor | (pH drop:stress metabolites)\|donor | *S. aurata* | CM - pH drop+SM | 0.8983 | 0.4092 | 2.1950 | 0.0563 |
| pH drop:stress metabolites:donor | (pH drop:stress metabolites)\|donor | *S. aurata* | pH drop - SM | -0.5224 | 0.5478 | -0.9536 | 0.4084 |
| pH drop:stress metabolites:donor | (pH drop:stress metabolites)\|donor | *S. aurata* | pH drop - pH drop+SM | -0.6391 | 0.5478 | -1.1666 | 0.3650 |
| pH drop:stress metabolites:donor | (pH drop:stress metabolites)\|donor | *S. aurata* | SM - pH drop+SM | -0.1167 | 0.4472 | -0.2609 | 0.7941 |
| pH drop:stress metabolites:donor | donor\|(pH drop: stress metabolites) | CM | *S. aurata - C. maenas* | -0.3678 | 0.3385 | -1.0865 | 0.3697 |
| pH drop:stress metabolites:donor | donor\|(pH drop: stress metabolites) | pH drop | *S. aurata - C. maenas* | 0.8806 | 0.5326 | 1.6535 | 0.1965 |
| pH drop:stress metabolites:donor | donor\|(pH drop: stress metabolites) | SM | *S. aurata - C. maenas* | 0.8425 | 0.3687 | 2.2850 | 0.0893 |
| pH drop:stress metabolites:donor | donor\|(pH drop: stress metabolites) | pH drop+SM | *S. aurata - C. maenas* | 0.2945 | 0.3843 | 0.7665 | 0.4434 |

**Table S9.** Results of post-hoc pairwise comparisons of interest from the interaction term between the three main predictors (pH, metabolites, donor) of the binomial generalised linear model on the avoidance behaviour (including freezing and escaping) in green shore crab (*Carcinus maenas*) receiving metabolites from conspecifics (*C. maenas*) or heterospecific gilt-head sea bream (*Sparus aurata*). Post-hoc tests are split into two terms expressed as '(pH drop:stress metabolites)|donor' and 'donor|(pH drop:stress metabolites)', respectively comparing the four treatments (CM, pH drop, SM, and pH drop+SM) within each donor group, and comparing the donor effect within each treatment (CM, pH drop, SM, or pH drop+SM). Significance ($P \leq 0.05$) is shown by p-values in bold. Results are expressed on the log (not the response) scale. SE: standard error of estimate. P-values are adjusted using false discovery rate adjustment for the number of comparisons within each post-hoc term. Comparisons were calculated using the emmeans R package (Lenth, 2019) on the generalised linear model computed via the stats R package (R Core Team, 2020). CM: control metabolites tested in control pH (pH = 8.2), pH drop: control metabolites tested in pH drop (pH = 7.6), SM: stress metabolites tested in control pH (pH = 8.2), pH drop+SM: stress metabolites tested in pH drop (pH = 7.6).

| **Model term** | **Post-hoc Term** | **Group** | **Comparison** | **Estimate** | **SE** | **Z** | **Padj** |
|---|---|---|---|---|---|---|---|
| pH drop:stress metabolites:donor | (pH drop:stress metabolites)\|donor | *C. maenas* | CM-pH drop | 1.1263 | 0.9173 | 1.2279 | 0.3292 |
| pH drop:stress metabolites:donor | (pH drop:stress metabolites)\|donor | *C. maenas* | CM-SM | 1.1263 | 0.9173 | 1.2279 | 0.3292 |
| pH drop:stress metabolites:donor | (pH drop:stress metabolites)\|donor | *C. maenas* | CM-pH drop+SM | 0.0000 | 0.7429 | 0.0000 | 1.0000 |
| pH drop:stress metabolites:donor | (pH drop:stress metabolites)\|donor | *C. maenas* | pH drop-SM | 0.0000 | 1.0628 | 0.0000 | 1.0000 |
| pH drop:stress metabolites:donor | (pH drop:stress metabolites)\|donor | *C. maenas* | pH drop-pH drop+SM | 1.1263 | 0.9173 | 1.2279 | 0.3292 |
| pH drop:stress metabolites:donor | (pH drop:stress metabolites)\|donor | *C. maenas* | SM-pH drop+SM | 1.1263 | 0.9173 | 1.2279 | 0.3292 |
| pH drop:stress metabolites:donor | (pH drop:stress metabolites)\|donor | *S. aurata* | CM-pH drop | -4.4288 | 1.2325 | **-3.5933** | **0.0020** |
| pH drop:stress metabolites:donor | (pH drop:stress metabolites)\|donor | *S. aurata* | CM-SM | -2.7536 | 0.8402 | **-3.2773** | **0.0031** |
| pH drop:stress metabolites:donor | (pH drop:stress metabolites)\|donor | *S. aurata* | CM-pH drop+SM | -1.8936 | 0.8141 | **-2.3259** | **0.0400** |
| pH drop:stress metabolites:donor | (pH drop:stress metabolites)\|donor | *S. aurata* | pH drop-SM | 1.6752 | 1.1639 | 1.4393 | 0.1801 |
| pH drop:stress metabolites:donor | (pH drop:stress metabolites)\|donor | *S. aurata* | pH drop-pH drop+SM | 2.5352 | 1.1465 | **2.2112** | **0.0405** |
| pH drop:stress metabolites:donor | (pH drop:stress metabolites)\|donor | *S. aurata* | SM-pH drop+SM | 0.8600 | 0.7148 | 1.2031 | 0.2289 |
| pH drop:stress metabolites:donor | donor\|(pH drop: stress metabolites) | CM | *S. aurata - C. maenas* | 0.4626 | 0.8326 | 0.5556 | 0.5785 |
| pH drop:stress metabolites:donor | donor\|(pH drop: stress metabolites) | pH drop | *S. aurata - C. maenas* | -5.0925 | 1.2919 | **-3.9417** | **0.0003** |
| pH drop:stress metabolites:donor | donor\|(pH drop: stress metabolites) | SM | *S. aurata - C. maenas* | -3.4173 | 0.9249 | **-3.6949** | **0.0004** |
| pH drop:stress metabolites:donor | donor\|(pH drop: stress metabolites) | pH drop+SM | *S. aurata - C. maenas* | -1.4310 | 0.7197 | -1.9882 | 0.0624 |

**Table S10**. Results of binomial generalised linear model for the main effects of predictors (pH drop, stress metabolites, donor) on the freezing behaviour in green shore crab (*Carcinus maenas*). Significance ($P \leq 0.05$) of is shown by p-values in bold. Sample size: 320 observations. Overall significance of models from Chi-squared analyses of deviance when including only predictors was P = 0.009. Covariates were dropped from the model after an analysis of deviance showed that they passed the Chi-squared test (number of water uses: P = 0.8469, crab size: P = 0.0798). SE is the standard error of estimate. Modelled using the stats R package (R Core Team, 2020).

| **Predictor** | **Estimate** | **SE** | **Z** | **P** |
|---|---|---|---|---|
| (Intercept) | -1.3863 | 0.5590 | -2.4799 | 0.0131 |
| pH drop | -0.8109 | 0.9317 | -0.8704 | 0.3841 |
| stress metabolites | -1.5581 | 1.1683 | -1.3336 | 0.1823 |
| donor | -0.6286 | 0.9376 | -0.6704 | 0.5026 |
| pH drop:stress metabolites | 2.6568 | 1.4789 | 1.7964 | 0.0724 |
| pH drop:donor | 2.8258 | 1.2980 | **2.1771** | **0.0295** |
| stress metabolites:donor | 2.7257 | 1.4730 | 1.8505 | 0.0642 |
| pH drop:stress metabolites:donor | -4.0475 | 1.8620 | **-2.1738** | **0.0297** |

**Table S11**. Results of post-hoc pairwise comparisons of interest from the interaction term between the three main predictors (pH drop, stress metabolites, donor) of the binomial generalised linear model on the freezing behaviour in green shore crab (*Carcinus maenas*) receiving metabolites from conspecifics (*C. maenas*) or heterospecific gilt-head sea bream (*Sparus aurata*). Post-hoc tests are split into two terms expressed as '(pH drop:stress metabolites)|donor' and 'donor|(pH drop:stress metabolites)', respectively comparing the four treatments (CM, pH drop, SM, and pH drop+SM) within each donor group, and comparing the donor effect within each treatment (CM, pH drop, SM, or pH drop+SM). Significance ($P \leq 0.05$) is shown by p-values in bold. Results are expressed on the log (not the response) scale. SE: standard error of estimate. P-values are adjusted using false discovery rate adjustment for the number of comparisons within each post-hoc term. Comparisons were calculated using the emmeans R package (Lenth, 2019) on the generalised linear model computed via the stats R package (R Core Team, 2020). CM: control metabolites tested in control pH (pH = 8.2), pH drop: control metabolites tested in pH drop (pH = 7.6), SM: stress metabolites tested in control pH (pH = 8.2), pH drop+SM: stress metabolites tested in pH drop (pH = 7.6).

| Model term | Post-hoc Term | Group | Comparison | Estimate | SE | Z | Padj |
|---|---|---|---|---|---|---|---|
| pH drop:stress metabolites:donor | (pH drop:stress metabolites)\|donor | *C. maenas* | CM-pH drop | 0.8109 | 0.9317 | 0.8704 | 0.5761 |
| pH drop:stress metabolites:donor | (pH drop:stress metabolites)\|donor | *C. maenas* | CM-SM | 1.5581 | 1.1683 | 1.3336 | 0.4514 |
| pH drop:stress metabolites:donor | (pH drop:stress metabolites)\|donor | *C. maenas* | CM-pH drop+SM | -0.2877 | 0.7610 | -0.3780 | 0.7054 |
| pH drop:stress metabolites:donor | (pH drop:stress metabolites)\|donor | *C. maenas* | pH drop-SM | 0.7472 | 1.2681 | 0.5892 | 0.6668 |
| pH drop:stress metabolites:donor | (pH drop:stress metabolites)\|donor | *C. maenas* | pH drop-pH drop+SM | -1.0986 | 0.9068 | -1.2116 | 0.4514 |
| pH drop:stress metabolites:donor | (pH drop:stress metabolites)\|donor | *C. maenas* | SM-pH drop+SM | -1.8458 | 1.1486 | -1.6074 | 0.4514 |
| pH drop:stress metabolites:donor | (pH drop:stress metabolites)\|donor | *S. aurata* | CM-pH drop | -2.0149 | 0.9037 | -2.2396 | 0.1321 |
| pH drop:stress metabolites:donor | (pH drop:stress metabolites)\|donor | *S. aurata* | CM-SM | -1.1676 | 0.8974 | -1.3016 | 0.3378 |
| pH drop:stress metabolites:donor | (pH drop:stress metabolites)\|donor | *S. aurata* | CM-pH drop+SM | -1.7918 | 0.8898 | -2.0138 | 0.1331 |
| pH drop:stress metabolites:donor | (pH drop:stress metabolites)\|donor | *S. aurata* | pH drop-SM | 0.8573 | 0.6986 | 1.2128 | 0.3378 |
| pH drop:stress metabolites:donor | (pH drop:stress metabolites)\|donor | *S. aurata* | pH drop-pH drop+SM | 0.2231 | 0.6892 | 0.3238 | 0.7461 |
| pH drop:stress metabolites:donor | (pH drop:stress metabolites)\|donor | *S. aurata* | SM-pH drop+SM | -0.6242 | 0.6805 | -0.9182 | 0.4309 |
| pH drop:stress metabolites:donor | donor\|(pH drop: stress metabolites) | CM | *S. aurata - C. maenas* | 0.6286 | 0.9376 | 0.6704 | 0.5026 |
| pH drop:stress metabolites:donor | donor\|(pH drop: stress metabolites) | pH drop | *S. aurata - C. maenas* | -2.1972 | 0.8975 | -2.4481 | 0.0574 |
| pH drop:stress metabolites:donor | donor\|(pH drop: stress metabolites) | SM | *S. aurata - C. maenas* | -2.0971 | 1.1361 | -1.8460 | 0.1298 |
| pH drop:stress metabolites:donor | donor\|(pH drop: stress metabolites) | pH drop+SM | *S. aurata - C. maenas* | -0.8755 | 0.7012 | -1.2485 | 0.2824 |

**Table S12**. Results of binomial generalised linear model for the main effects of predictors (pH drop, stress metabolites, donor) on the escaping behaviour in green shore crab (*Carcinus maenas*). Significance ($P \leq 0.05$) is shown by p-values in bold. Sample size: 151 observations. Overall significance of models from Chi-squared analyses of deviance when including only predictors was $P < 0.0001$. Covariates were dropped from the model after an analysis of deviance showed that they passed the Chi-squared test (number of water uses: $P = 0.2393$, crab size: $P = 0.3273$). SE is the standard error of estimate. Modelled using the stats R package (R Core Team, 2020).

| Predictor | Estimate | SE | Z | P |
|---|---|---|---|---|
| (Intercept) | -2.1972 | 0.7454 | -2.9479 | 0.0032 |
| pH drop | -17.3688 | 2404.6705 | -0.0072 | 0.9942 |
| stress metabolites | 0.0000 | 1.0541 | 0.0000 | 1.0000 |
| donor | -0.5754 | 1.2720 | -0.4523 | 0.6510 |
| pH drop:stress metabolites | 0.0000 | 3400.7177 | 0.0000 | 1.0000 |
| pH drop:donor | 21.6078 | 2404.6708 | 0.0090 | 0.9928 |
| stress metabolites:donor | 3.1781 | 1.5434 | **2.0592** | **0.0395** |
| pH drop:stress metabolites:donor | -5.5999 | 3400.7179 | -0.0016 | 0.9987 |

**Table S13**. Results of post-hoc pairwise comparisons of interest from the interaction term between the three main predictors (pH drop, stress metabolites, donor) of the binomial generalised linear model on the escaping behaviour in green shore crab (*Carcinus maenas*) receiving metabolites from conspecifics (*C. maenas*) or heterospecific gilt-head sea bream (*Sparus aurata*). Post-hoc tests are split into two terms expressed as '(pH drop:stress metabolites)|donor' and 'donor|(pH drop:stress metabolites)', respectively comparing the four treatments (CM, pH drop, SM, and pH drop+SM) within each donor group, and comparing the donor effect within each treatment (CM, pH drop, SM, or pH drop+SM). Significance ($P \leq 0.05$) is shown by p-values in bold. Results are expressed on the log (not the response) scale. SE: standard error of estimate. P-values are adjusted using false discovery rate adjustment for the number of comparisons within each post-hoc term. Comparisons were calculated using the emmeans R package (Lenth, 2019) on the generalised linear model computed via the stats R package (R Core Team, 2020). CM: control metabolites tested in control pH (pH = 8.2), pH drop: control metabolites tested in pH drop (pH = 7.6), SM: stress metabolites tested in control pH (pH = 8.2), pH drop+SM: stress metabolites tested in pH drop (pH = 7.6).

| **Model term** | **Post-hoc Term** | **Subset** | **Comparison** | **Estimate** | **SE** | **Z** | **Padj** |
|---|---|---|---|---|---|---|---|
| pH drop:stress metabolites:donor | (pH drop:stress metabolites)\|donor | *C. maenas* | CM - pH drop | 17.3688 | 2404.6705 | 0.0072 | 1.0000 |
| pH drop:stress metabolites:donor | (pH drop:stress metabolites)\|donor | *C. maenas* | CM - SM | 0.0000 | 1.0541 | 0.0000 | 1.0000 |
| pH drop:stress metabolites:donor | (pH drop:stress metabolites)\|donor | *C. maenas* | CM - pH drop+SM | 17.3688 | 2404.6705 | 0.0072 | 1.0000 |
| pH drop:stress metabolites:donor | (pH drop:stress metabolites)\|donor | *C. maenas* | pH drop - SM | -17.3688 | 2404.6705 | -0.0072 | 1.0000 |
| pH drop:stress metabolites:donor | (pH drop:stress metabolites)\|donor | *C. maenas* | pH drop - pH drop+SM | 0.0000 | 3400.7175 | 0.0000 | 1.0000 |
| pH drop:stress metabolites:donor | (pH drop:stress metabolites)\|donor | *C. maenas* | SM - pH drop+SM | 17.3688 | 2404.6705 | 0.0072 | 1.0000 |
| pH drop:stress metabolites:donor | (pH drop:stress metabolites)\|donor | *S. aurata* | CM - pH drop | -4.2389 | 1.2136 | **-3.4929** | **0.0029** |
| pH drop:stress metabolites:donor | (pH drop:stress metabolites)\|donor | *S. aurata* | CM - SM | -3.1781 | 1.1273 | **-2.8191** | **0.0096** |
| pH drop:stress metabolites:donor | (pH drop:stress metabolites)\|donor | *S. aurata* | CM - pH drop+SM | -1.8171 | 1.1573 | -1.5701 | 0.1397 |
| pH drop:stress metabolites:donor | (pH drop:stress metabolites)\|donor | *S. aurata* | pH drop - SM | 1.0609 | 0.7865 | 1.3488 | 0.1774 |
| pH drop:stress metabolites:donor | (pH drop:stress metabolites)\|donor | *S. aurata* | pH drop - pH drop+SM | 2.4218 | 0.8290 | **2.9215** | **0.0096** |
| pH drop:stress metabolites:donor | (pH drop:stress metabolites)\|donor | *S. aurata* | SM - pH drop+SM | 1.3610 | 0.6966 | 1.9537 | 0.0761 |
| pH drop:stress metabolites:donor | donor\|(pH drop: stress metabolites) | CM | *S. aurata - C. maenas* | 0.5754 | 1.2720 | 0.4523 | 0.9938 |
| pH drop:stress metabolites:donor | donor\|(pH drop: stress metabolites) | pH drop | *S. aurata - C. maenas* | -21.0324 | 2404.6710 | -0.0087 | 0.9938 |
| pH drop:stress metabolites:donor | donor\|(pH drop: stress metabolites) | SM | *S. aurata - C. maenas* | -2.6027 | 0.8740 | **-2.9779** | **0.0116** |
| pH drop:stress metabolites:donor | donor\|(pH drop: stress metabolites) | pH drop+SM | *S. aurata - C. maenas* | -18.6106 | 2404.6710 | -0.0077 | 0.9938 |

**Table S14**. Results of two-way ANOVA for the main effects of pH drop and stress metabolites on the respiration rates in green shore crab (*Carcinus maenas*) in presence of metabolites from conspecifics. Overall significance of models from Chi-squared analyses of deviance was P = 0.1446. Significance ($P \leq 0.05$) is shown by p-values in bold. Sample size: 151 observations. SE is the standard error of estimate. Effect size thresholds followed the classification given in Sawilowsky (2009). Post-hoc tests were all nonsignificant.

| **Predictor** | **Sum Square** | **% explained variance** | **F** | **P** | **Cohen's \|d\|** |
|---|---|---|---|---|---|
| pH drop | 0.4258 | 0.5390 | 0.4391 | 0.5095 | \|d\| = 0.15, '*very small*' |
| stress metabolites | 1.1376 | 1.4400 | 1.1732 | 0.2822 | \|d\| = 0.24, '*small'* |
| pH drop:stress metabolites | 3.7439 | 4.7392 | **3.8612** | **0.0531** | **\|d\| = 0.68, *'medium'*** |
| Residuals | 73.6927 | 93.2818 | - | - | - |

**Table S15.** Results of post-hoc pairwise comparisons of interest from the interaction term between the three main predictors (pH, metabolites, donor) of the Cox proportional hazard model on the time-to-success analysis in harbour ragworm (*Hediste diversicolor*) receiving metabolites from conspecifics (*H. diversicolor*) or heterospecific gilt-head sea bream (*Sparus aurata*). Post-hoc tests are split into two terms expressed as '(pH drop:stress metabolites)|donor' and 'donor|(pH drop:stress metabolites)', respectively comparing the four treatments (CM, pH drop, SM, and pH drop+SM) within each donor group, and comparing the donor effect within each treatment (CM, pH drop, SM, or pH drop+SM). Events are the success to completely burrow the head in the sand in 300 seconds. Significance ($P \leq 0.05$) is shown by p-values in bold. Results are expressed on the log (not the response) scale. SE: standard error of estimate. P-values are adjusted using false discovery rate adjustment for the number of comparisons within each post-hoc term. Comparisons were calculated using the emmeans R package (Lenth, 2019) on the Cox proportional hazard model computed via the survival R package (Therneau and Grambsch, 2000; Therneau, 2020). CM: control metabolites tested in control pH (pH = 8.2), pH drop: control metabolites tested in pH drop (pH = 7.6), SM: stress metabolites tested in control pH (pH = 8.2), pH drop+SM: stress metabolites tested in pH drop (pH = 7.6).

| **Interaction term** | **Post-hoc Term** | **Group** | **Comparison** | **Estimate** | **SE** | **Z** | **Padj** |
|---|---|---|---|---|---|---|---|
| pH drop:stress metabolites:donor | (pH drop:stress metabolites)\|donor | *H. diversicolor* | CM - pH drop | 1.1899 | 0.3045 | **3.9075** | **0.0006** |
| pH drop:stress metabolites:donor | (pH drop:stress metabolites)\|donor | *H. diversicolor* | CM - SM | 1.0442 | 0.3049 | **3.4251** | **0.0012** |
| pH drop:stress metabolites:donor | (pH drop:stress metabolites)\|donor | *H. diversicolor* | CM - pH drop+SM | 0.8500 | 0.2453 | **3.4652** | **0.0012** |
| pH drop:stress metabolites:donor | (pH drop:stress metabolites)\|donor | *H. diversicolor* | pH drop - SM | -0.1456 | 0.3652 | -0.3988 | 0.6900 |
| pH drop:stress metabolites:donor | (pH drop:stress metabolites)\|donor | *H. diversicolor* | pH drop - pH drop+SM | -0.3399 | 0.3181 | -1.0684 | 0.4280 |
| pH drop:stress metabolites:donor | (pH drop:stress metabolites)\|donor | *H. diversicolor* | SM - pH drop+SM | -0.1943 | 0.3182 | -0.6104 | 0.6499 |
| pH drop:stress metabolites:donor | (pH drop:stress metabolites)\|donor | *S. aurata* | CM - pH drop | 0.2869 | 0.2315 | 1.2392 | 0.2583 |
| pH drop:stress metabolites:donor | (pH drop:stress metabolites)\|donor | *S. aurata* | CM - SM | 0.4951 | 0.2317 | 2.1365 | 0.0653 |
| pH drop:stress metabolites:donor | (pH drop:stress metabolites)\|donor | *S. aurata* | CM - pH drop+SM | 0.8714 | 0.2605 | **3.3452** | **0.0049** |
| pH drop:stress metabolites:donor | (pH drop:stress metabolites)\|donor | *S. aurata* | pH drop - SM | 0.2082 | 0.2358 | 0.8828 | 0.3773 |
| pH drop:stress metabolites:donor | (pH drop:stress metabolites)\|donor | *S. aurata* | pH drop- pH drop+SM | 0.5845 | 0.2639 | 2.2151 | 0.0653 |
| pH drop:stress metabolites:donor | (pH drop:stress metabolites)\|donor | *S. aurata* | SM - pH drop+SM | 0.3763 | 0.2639 | 1.4261 | 0.2308 |
| pH drop:stress metabolites:donor | donor\|(pH drop: stress metabolites) | CM | *S. aurata - H. diversicolor* | 0.3616 | 0.2258 | 1.6013 | 0.2186 |
| pH drop:stress metabolites:donor | donor\|(pH drop: stress metabolites) | pH drop | *S. aurata - H. diversicolor* | -0.5413 | 0.3076 | -1.7598 | 0.2186 |
| pH drop:stress metabolites:donor | donor\|(pH drop: stress metabolites) | SM | *S. aurata - H. diversicolor* | -0.1875 | 0.3079 | -0.6090 | 0.5425 |
| pH drop:stress metabolites:donor | donor\|(pH drop: stress metabolites) | pH drop+SM | *S. aurata - H. diversicolor* | 0.3831 | 0.2761 | 1.3876 | 0.2204 |

**Table S16**. Results of post-hoc pairwise comparisons of interest from the interaction term between the main predictors (pH drop, stress metabolites) of the binomial generalised linear model on the avoidance behaviour (including freezing, curling, flipping, and slime secretion) in harbour ragworm (*Hediste diversicolor*) receiving metabolites from conspecifics (*H. diversicolor*) or heterospecific gilt-head sea bream (*Sparus aurata*). Due to missing observations in conspecific control metabolites at control pH (CM) treatment, the model was built only for treatments SM, pH drop, and pH drop+SM allowing for post-hoc comparisons between those three treatments. In the S. aurata group, the model included the interactive effects of pH drop and stress metabolites allowing for post-hoc comparisons between the four treatments (CM, SM, pH drop, pH drop+SM). Significance ($P \leq 0.05$) is shown by p-values in bold. Results are expressed on the log (not the response) scale. SE: standard error of estimate. P-values are adjusted using false discovery rate adjustment for the number of comparisons within each post-hoc term. Comparisons were calculated using the emmeans R package (Lenth, 2019) on the generalised linear model computed via the stats R package (R Core Team, 2020). CM: control metabolites tested in control pH (pH = 8.2), pH drop: control metabolites tested in pH drop (pH = 7.6), SM: stress metabolites tested in control pH (pH = 8.2), pH drop+SM: stress metabolites tested in pH drop (pH = 7.6).

| **Post-hoc Term** | **Group** | **Comparison** | **Estimate** | **SE** | **Z** | **Padj** |
|---|---|---|---|---|---|---|
| Treatment | *H. diversicolor* | pH drop - pH_drop+SM | 18.8729 | 1603.1137 | 0.0118 | 0.9906 |
| Treatment | *H. diversicolor* | pH drop - SM | -0.3185 | 0.5658 | -0.5629 | 0.9906 |
| Treatment | *H. diversicolor* | pH drop+SM - SM | -19.1914 | 1603.1137 | -0.0120 | 0.9906 |
| pH drop:stress metabolites | *S. aurata* | CM - pH drop | 18.8729 | 1603.1137 | 0.0120 | 0.9906 |
| pH drop:stress metabolites | *S. aurata* | CM - SM | -3.3557 | 0.7788 | **-4.3086** | **0.0001** |
| pH drop:stress metabolites | *S. aurata* | CM - pH drop+SM | -0.7376 | 0.5822 | -1.2670 | 0.4103 |
| pH drop:stress metabolites | *S. aurata* | pH drop - SM | -22.2287 | 1603.1137 | -0.0139 | 0.9906 |
| pH drop:stress metabolites | *S. aurata* | pH drop - pH drop+SM | -19.6105 | 1603.1136 | -0.0122 | 0.9906 |
| pH drop:stress metabolites | *S. aurata* | SM - pH drop+SM | 2.6181 | 0.6675 | **3.9225** | **0.0003** |

**Table S17**. Results of post-hoc pairwise comparisons from Cox proportional hazard models on the time-to-success analysis in small hermit crab (*Diogenes pugilator*) (model 1) and harbour ragworm (*Hediste diversicolor*) (model 2) receiving metabolites from heterospecific gilt-head sea bream (*Sparus aurata*). Events are the success to find a feeding cue in *D. pugilator* or to burrow the head in *H. diversicolor* in 300 seconds. Unsuccessful animals are censored. Significance ($P \leq 0.05$) is shown by p-values in bold. Overall significance of models from Chi-squared analyses of deviance were: *D. pugilator*: P = 0.8503; *H. diversicolor*: P = 0.0137. Covariates were dropped from the model after an analysis of deviance showed that they passed the Chi-squared test both in model 1 (number of water uses: P = 0.4551, crab size: P = 0.7333) and model 2 (number of water uses: P = 0.7566). Results are expressed on the log (not the response) scale. SE: standard error of estimate. P-values are adjusted using false discovery rate adjustment for the number of pairwise comparisons in each species. Comparisons were calculated using the emmeans R package (Lenth, 2019) on the Cox proportional hazard model computed via the survival R package (Therneau and Grambsch, 2000; Therneau, 2020). CM100%: undiluted control metabolites tested in control pH (pH = 8.2), CM: tenfold diluted control metabolites tested in control pH (pH = 8.2), SM: tenfold diluted stress metabolites tested in control pH (pH = 8.2).

| Species | Comparison | Success ratio | Estimate | SE | Z | Padj |
|---|---|---|---|---|---|---|
| *D. pugilator* | CM-CM100% | 0.97 | 0.0285 | 0.4472 | 0.0638 | 0.9492 |
| | CM-SM | 0.97 | -0.1943 | 0.4282 | -0.4538 | 0.9492 |
| | CM100%-SM | 1.21 | -0.2228 | 0.4282 | -0.5203 | 0.9492 |
| *H. diversicolor* | CM-CM100% | 1.04 | -0.0407 | 0.2244 | -0.1812 | 0.8562 |
| | CM-SM | 1.04 | 0.5511 | 0.2314 | **2.3714** | **0.0266** |
| | CM100%-SM | 0.58 | 0.5917 | 0.2291 | **2.5828** | **0.0266** |

**Table S18**. Results of post-hoc pairwise comparisons from binomial generalised linear models on the escaping behaviour (model 1) in small hermit crab (*Diogenes pugilator*) and the avoidance behaviour (including freezing, curling, flipping, and slime secretion) (model 2) in harbour ragworm (*Hediste diversicolor*) receiving metabolites from heterospecific gilt-head sea bream (*Sparus aurata*). Significance ($P \leq 0.05$) is shown by p-values in bold. Overall significance of models from Chi-squared analyses of deviance were: *D. pugilator*: P = 0.0078; *H. diversicolor*: P < 0.0001. Covariates were dropped from the model after an analysis of deviance showed that they passed the Chi-squared test both in model 1 (number of water uses: P = 0.5274, crab size: P = 0.0983) and model 2 (number of water uses: P = 0.4201). Results are expressed on the log (not the response) scale. SE: standard error of estimate. P-values are adjusted using false discovery rate adjustment for the number of pairwise comparisons in each species. Comparisons were calculated using the emmeans R package (Lenth, 2019) on the generalised linear model computed via the stats R package (R Core Team, 2020). CM100%: undiluted control metabolites tested in control pH (pH = 8.2), CM: tenfold diluted control metabolites tested in control pH (pH = 8.2), SM: tenfold diluted stress metabolites tested in control pH (pH = 8.2).

| Species | Comparison | Estimate | SE | Z | Padj |
|---|---|---|---|---|---|
| *D. pugilator* | CM-CM100% | 0.0000 | 1.0260 | 0.0000 | 1.0000 |
| | CM-SM | -1.8458 | 0.8122 | **-2.2728** | **0.0346** |
| | CM100%-SM | -1.8458 | 0.8122 | **-2.2728** | **0.0346** |
| *H. diversicolor* | CM-CM100% | 0.8383 | 0.6341 | 1.3222 | 0.1861 |
| | CM-SM | -3.3557 | 0.7787 | **-4.3087** | **< 0.0001** |
| | CM100%-SM | -4.1941 | 0.7132 | **-5.8809** | **< 0.0001** |
